# Apparent cooperativity between human CMV virions introduces errors in conventional methods of calculating multiplicity of infection

**DOI:** 10.1101/2025.04.23.650360

**Authors:** Christopher Peterson, Joshua Miller, Brent J. Ryckman, Vitaly V. Ganusov

## Abstract

Whether infection of cells by individual virions occurs randomly or if there is some form(s) of competition or cooperativity between individual virions remains largely unknown for most virus-cell associations. Here we studied cooperativity/competition for three different strains of human cytomegalovirus (**HCMV**) on two different cell types (fibroblasts and epithelial cells). By titrating viral inocula concentrations in small steps over several orders of magnitude, and by using flow cytometry to precisely measure frequency of infected cells, we found that for most virus-cell associations, the frequency of cell infection increases faster than linear with an increasing inoculum concentration, indicating cooperativity between individual infecting virions. Mathematical modeling suggests that this apparent cooperativity cannot be explained by heterogeneity in either the infectivity of the individual virions or the resistance of individual cells to infection, or by simple aggregation/clumping of viral particles. Stochastic simulations of two additional alternative models that allow for i) reduction in cell resistance to infection when exposed to multiple virions, or ii) compensation in infectivity of poorly infectious virions when coinfecting cells with more infectious virions, resulted in apparent viral cooperativity. Analysis of other published datasets suggests presence of apparent viral cooperativity for HIV and vaccinia virus, infecting CRFK or HeLa cells, respectively, but not for tobacco mosaic virus forming plaques on plant leaves. We thus 1) propose a methodology to rigorously evaluate apparent cooperativity of viruses infecting target cells, and 2) demonstrate that knowing the degree of virus cooperativity for any given virus-cell combination is important for an accurate quantification of multiplicity of infection (**MOI**).

**Graphical abstract:** 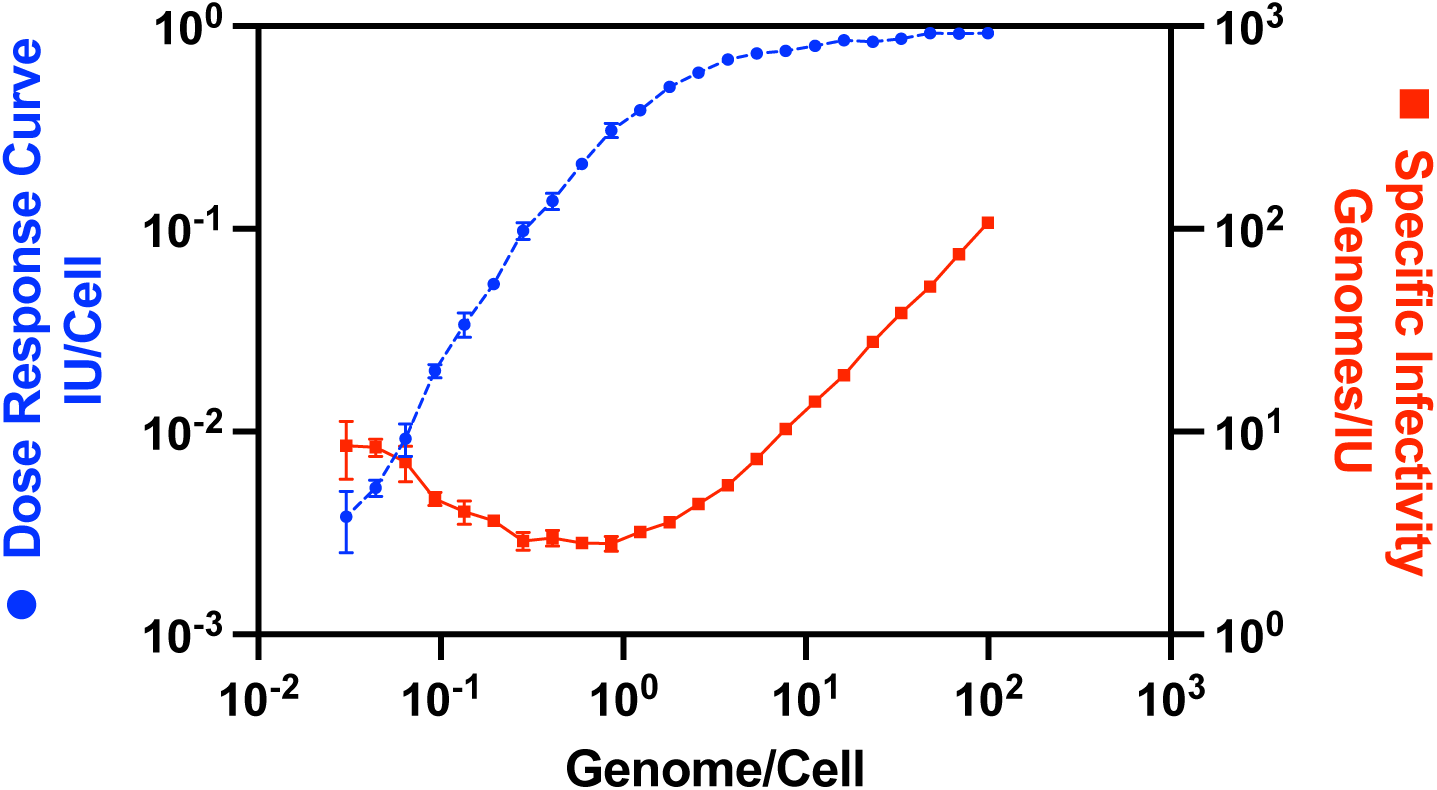

Infection Unit per cell (IU/Cell) and specific infectivity (Genome/IU) of human cytomegalovirus depend on the infecting dose (Genome/Cell). The data are for HCMV-TB infecting fibroblasts.

## Introduction

The infectivity of virus populations (i.e., virus stocks) is often reported as some value of Infectious Units (**IUs**), such as plaque-forming units (**PFUs**), per unit volume [1]. Such stock infectivity alone offers no insight into the potential of any individual virion to establish an infection, but instead reflects the gross phenotype of the entire virus population. Specific infectivity (**SI**), a ratio of the number of virions to IUs in a sample, better describes infection ability of individual virions even though it still reflects properties of the average virion. Actual SI values vary widely for different virus-cell combinations. For example, SI of Ebola virus was estimated to be around 500 particles to PFU while for varicella-zoster virus, 40,000 particles to PFU has been reported [2, 3]. Wide variation can also be observed among different strains within the same viral species and are further dependent on the target cell type. For example we reported particles to IU for three strains of human cytomegalovirus (**HCMV**) at 11 for strain TB40e/BAC4 (**TB**), 217 for TR, and between 5,825 and 100,000 for Merlin (**ME**) when IUs were measured on fibroblasts, but 419, 352,330 and 10,529, respectively, when determined on epithelial cells (**ECs**) [4, 5]. Although it is generally accepted that there are viral and cellular components responsible for varying SI values, the mechanisms responsible are poorly understood. Moreover, whether particle:PFU ratios *>* 1 reflect the number of non-infectious or otherwise defective viral particles in the sample or is instead a requirement for more than one virion to establish a single infection (i.e., virion cooperation) remains largely unknown.

When it is assumed that a single virion can establish infection and the target cells are not limiting, it is expected that dilution of the stock will result in a proportional decrease in the frequency of productive infections [6–8]. Such a linear relationship between the number of viral particles per target cell and the frequency of cell infection is expected from a single-hit model [8–10] and is used to then perform experiments with a desired degree of cellular infection as multiplicity of infection (**MOI**). Indeed, classical experiments showed a linear relationship between the concentration of viral stock and number of infections (e.g., in a plaque assay) [6, 7, 11]; however, deviations from the simple, single hit model have also been observed, especially in plant viruses [7, 10]. Following work on the probability of infection of a host when exposed to variable doses of pathogens, if target cells vary in their susceptibility to infection, it is expected that the frequency of cellular infection will scale sublinearly with higher viral concentrations [12–14]. In such a scenario, infection rises faster initially as highly susceptible cells are infected, but saturates at higher viral concentrations due to the difficulty of infecting the more resistant cells. Moreover, at high virus concentrations, target cells become limiting and there may even be interference between viral particles in infecting cells, including with defective interfering particles (**DIPs**) – a so-called Von Magnus phenomenon [15]. Because the frequency of cellular infection by viruses has typically been studied in plaque assays, which rely on the assumptions of single-hit kinetics, deviations of the infection rate from that predicted by the single hit model have been difficult to assess at lower virion per cell frequencies, and so it is typically assumed that at low MOI, infection rate scales linearly with the number of virions cells are exposed to [16]. However, if scaling of the cell infection frequency with the number of virions per cell is non-linear, experiments with MOI different from those used to titrate the viral stock may lead to fewer infected cells than anticipated and then to misleading results.

In this study we have used HCMV to systematically study how the frequency of infection of target cells relates to the number of virions present in the culture. HCMV is an enveloped, double-stranded DNA virus in the herpesvirus family with a prevalence rate from 40% to 100%, depending on the particular human population [17, 18]. Infection of immunocompromized individuals, such as people with AIDS, organ transplant recipients, or newborn infants can result in severe disease [19, 20] and infection of otherwise healthy immunocompetent adults may contribute to increased risk of aging related conditions such as Alzheimer’s disease [21, 22]. To track infection rates we used three different HCMV strains TB, TR, and ME each expressing GFP or mCherry in place of the US11 gene [23] and counted infected cells with flow cytometry, capable of detecting very low frequencies of infected cells (with the limit of detection LOD ≈ 0.01%). We also used a carefully designed strategy to dilute the viral stock with a low dilution factor (*d_f_* = 1.3 − 1.5) and used two types of target cells — fibroblasts that are highly susceptible to HCMV infection and are a typical cell type used in experiments with HCMV and epithelial cells that are notably less susceptible to infection [24]. Interestingly, we found that for most virus strain-host cell combinations, increase in the number of virions per target cell leads to a nonlinear change in the frequency of infected cells, and for most tested combinations, cell infection frequency increased faster than linearly (slope *n >* 1 on log-log scale) with increasing numbers of virions per cell at low virion/cell numbers, a phenomenon we defined as “apparent cooperativity”. This manifests in our data as fewer than expected infections as the viral stock is increasingly diluted, presumably because virions cooperate to infect a single cell, rather than each infecting on their own. By simulating the process of cellular infection we show that apparent cooperativity cannot arise with random virus-cell interactions, and in fact, random interactions produce results predicted by the single hit model. Also, the non-linear response is not due to variability of virion infectivity or differences in target cell susceptibility to infection. We developed three alternative hypotheses to explain the observed non-linear response and simulated those interactions. Faster than linear increase in the frequency of cellular infection with increasing numbers of virions/cell could be achieved if infecting virions compensate each other’s infectivity (viral compensation hypothesis), or if a cell loses its ability to resist infection when exposed to multiple virions (accrued damage hypothesis), or, under some parameter values, when virions aggregate into clumps (aggregation hypothesis). Importantly, we show that apparent cooperativity will bias calculated MOI in experiments where infectivity of the viral stock was measured at a different MOI than that later used for experimentation. Our results also suggest that attributing specific infectivity to viral strains as an immutable trait is fundamentally incorrect as it is as much a property of target cells as it is of the virus or strain, and so references listing particle to PFU ratios should be updated to also include target cell-type and the number of virions/cell used in experiments to calculate the listed specific infectivity.

## Materials and methods

### Experimental data

#### Cell lines

Human foreskin fibroblast cells (**HFFCs** or fibroblasts) and MRC5 cells (also fibroblasts) were cultured in Dulbecco’s modified Eagle’s medium (**DMEM**, Sigma) supplemented with 5% heat-inactivated fetal bovine serum (**FBS**, Rocky Mountain Biologicals, Missoula, MT, USA) and 5% Fetalgro® (Rocky Mountain Biologicals, Missoula, MT, USA). We used MRC5 cells in the first step of recovering infection HCMV from BAC DNA (electroporation). For main experiments we used HFFCs. Human retinal pigment epithelial cells (**ECs** or ARPE-19, American Type Culture Collection, Manassas, VA, USA) were cultured in a 1:1 mixture of DMEM and Ham’s F-12 medium (DMEM:F-12, Gibco) and supplemented with 10% FBS.

#### Virus stocks

All HCMV strains were derived from bacterial artificial chromosome (BAC) clones. BAC clone TB40/e (BAC4, dubbed **TB**) was provided by Christian Sinzger (University of Ulm, Germany) [25]. BAC clone **TR** was provided by Jay Nelson (Oregon Health and Sciences University, Portland, OR, USA). BAC clone Merlin or **ME** (pAL1393), which contains tetracycline operator sequences within the transcriptional promoter of UL130 and UL131, was provided by Richard Stanton (Cardiff University, Cardiff, United Kingdom) [26]. All BAC clones were modified to express green fluorescent protein (**GFP**) or the monomeric red fluorescent protein mCherry (**mCherry**) with En passant recombineering by replacing US11 with the eGFP or mCherry gene, respectively. US11 is a resident ER protein that is considered an “immune evasion factor”. It promotes ERAD of MHC I and has no observable effect on replication of HCMV in cultured cells [27, 28]. Infectious HCMV was recovered by electroporation of BAC-DNA into MRC5 cells which were then co-cultured with either HFFCs (TB and TR) or HFF-tet cells (ME).

#### Quantification of viral stock infectivity using standard 10-fold dilutions

Confluent 6-well plates of HFFCs (or neonatal human dermal fibroblasts (**NHDF**) were inoculated with 1.0 mL/well of GFP or mCherry-expressing recombinant virus stock, in technical duplicates, at 1:10, 1:100, and 1:1000 dilutions. After a four-hour incubation at 37C, the inoculum was aspirated and replaced with 3.0 mL 2% FBS DMEM. At 3 days post-infection, cells were rinsed with PBS, lifted with trypsin, and quenched with 10% FBS DMEM before counting with a hemocytometer. Cells were then pelleted by centrifuging at 500g for 5 minutes and fixed with 4% paraformaldehyde for 20 minutes at room temperature (**RT**) after aspirating media. Cells were then pelleted again at 500g and resuspended in PBS. GFP-positive cells were counted using a Life Technologies Attune NxT Acoustic Focusing Cytometer and analyzed in FlowJo v10.

#### Virus Stock Dilution and Dose-Response Assay

Typically, serial dilutions are performed in 10-fold steps, allowing for coverage of a wide range of concentrations to increase the chance that at least one of those dilutions will result in an easily countable number of infections. We found it necessary to perform very small-fold dilutions (i.e., *d_f_* = 1.3 − 1.5) to increase the resolution of our dose-response curves. We limited our total number of dilutions to 23, plus a mock-infection control, so that a single strain-cell type combination could be assayed with three 24-well plates, allowing for technical triplicates of each concentration. The dilution factor was tailored for each strain-cell type pair to ensure that all the aspects of the dose-response curve could be analyzed, including the highest dilutions, where infections were undetectable, through the analytical region, and including saturation. If a too-small dilution factor were used, too many of the data points would reside within the saturation region of the curve, and the lower end of the curve would not be probed. This resulted in the dilution factor in our experiments to be *d_f_* = 1.3 − 1.5. The dilution factor was necessarily higher for strains with higher cell-free infectivity (e.g., TB) and also when using highly susceptible cells (e.g., fibroblasts). This allows the dose-response curve to quickly pass through the saturation portion and allow for complete coverage. Conversely, a strain with low infectivity or an assay on cells resistant to infection requires a smaller dilution factor, otherwise, the dilution series would quickly pass through the analytical region and not provide enough resolution for quality results.

We developed a methodology to allow for a consistent, low-dilution factor series, without the requirement to adjust pipette volumes during the experiment. We begin with a large volume of undiluted virus stock, typically 2 to 4 mL, and perform the dilutions as part of the inoculation process. We chose two 1000 uL pipettes and set one to 250 uL, the inoculation volume per well, and the other to 750 uL, the total inoculation volume for triplicate wells. We then measured the mass, after wetting the pipette tip, of 20^0^C DMEM (2%FBS) dispensed by each pipette three times to 0.001 grams. These masses were then converted to volume using the standard curve (**Supplemental Table S1** and **Supplemental Figure S1**). Three 24-well tissue culture plates were seeded at about 2 × 10^4^ cells per well with fibroblasts (HFFCs) or about 6 × 10^4^ cells per well with ECs and allowed to grow to confluency. We aspirated media from triplicate wells, rinsed with PBS, and aspirated again just prior to inoculation.

Starting with undiluted virus stock, triplicate wells were inoculated, using the pipette set to 250 uL after wetting the pipette tip. Then, using the pipette set to 750 uL, 20^0^C DMEM (2% FBS) was added to the virus stock to replace the volume just removed and the stock was gently vortexed to mix. This diluted stock was then used to inoculate the next set of triplicate wells and the process was repeated through the 23rd set of wells. The last set of wells were mock inoculated as a control. The plates were then incubated at 37^0^C for 4 hours before aspirating the inoculum and replacing it with 20^0^C DMEM (2% FBS). The plates continued incubation at 37^0^C for three days before the cells were fixed for analysis.

As the working stock is always diluted by the same volume, using the 750 uL pipette, the starting volume dictates the dilution factor. A larger starting volume is diluted, proportionally, less than a smaller starting volume. We developed this dilution method to reduce the potential for pipetting error, first by using only two pipettes, whose volumes are not adjusted during the experiment, secondly by producing a standard curve to calculate the volumes dispensed (**Supplemental Table S1**), and thirdly by performing the dilutions in a single conical tube, thereby decreasing the chance of volume inaccuracy caused by pipetting between containers.

While the dose-response assay would yield valid results by reporting the data using relative concentrations, we chose to anchor the x-axis by using the same units as the resulting infection rates, reporting concentrations as virus particles (genomes) per cell. We determined the concentration of the undiluted virus stock by isolating viral DNA from 200 uL of stock and utilizing qPCR (see below), to count and then calculate the number of viral genomes per 1 mL of stock. Using the pipette volumes determined with our standard curve, we could then calculate the viral genomes present in the inoculant in a well at each concentration.

#### DMEM Volume Standard Curve

DMEM (2% FBS) at 20^0^C was dispensed using a 1.0 ml serological pipette (FisherBrand cat num 13-678-11B, 2% tolerance) in triplicates at 1000, 900, 800, 750, 700, 600, 500, 400, 300, 250, 200, and 100 ul volumes and weighed to 0.0001 grams. This data was used to create a standard curve within GraphPad Prism and the dose-response assay volumes were determined by interpolation using that software.

#### Flow cytometry

We performed preliminary experiments to determine that measuring the frequency of infected cells after first round of infection is best done at three days after exposure of cells to the virus (**Supplemental Figures S2 and S3**). Three days post infection cells were washed with PBS and lifted with trypsin. Trypsin was quenched with DMEM containing 10% FBS and cells were spun at 500g for 5 min at RT. Cells were fixed with 4% paraformaldehyde (in PBS) for 15 min at RT before spinning as before and aspirating the fixing buffer. Cells were resuspended in 1 mL PBS and analyzed using an Attune NxT flow cytometer. Cells were separated from debris using FSC-A and SSC-A to gate, and single cells were gated using FSC-W and FSC-H. GFP+ and mCherry+ gates were drawn using mock-infected cells as negative controls.

#### qPCR

To quantify the viral particle/genome concentration in each stock, a real-time quantitative PCR (qPCR) assay was used to quantify viral DNA molecules, performed as previously described (Zhou 2015). A PureLink Viral RNA/DNA mini kit (Thermo Scientific) was used to isolate viral DNA and a sequence within UL83, conserved among ME, TR, and TB, was chosen as the amplicon. Viral genomes were quantified by SYBR green qPCR. Standard curves were performed using serial dilutions of a single PCR DNA product containing the sequence of the viral UL83. Sample Cq values were fitted to the standard curve to determine viral genomes per 1.0 mL of stock and reported as viral genomes per mL. We then used the value of genomes per mL and known concentration of target cell per mL to calculate genomes/cell (**Figure 1**).

**Figure 1:**
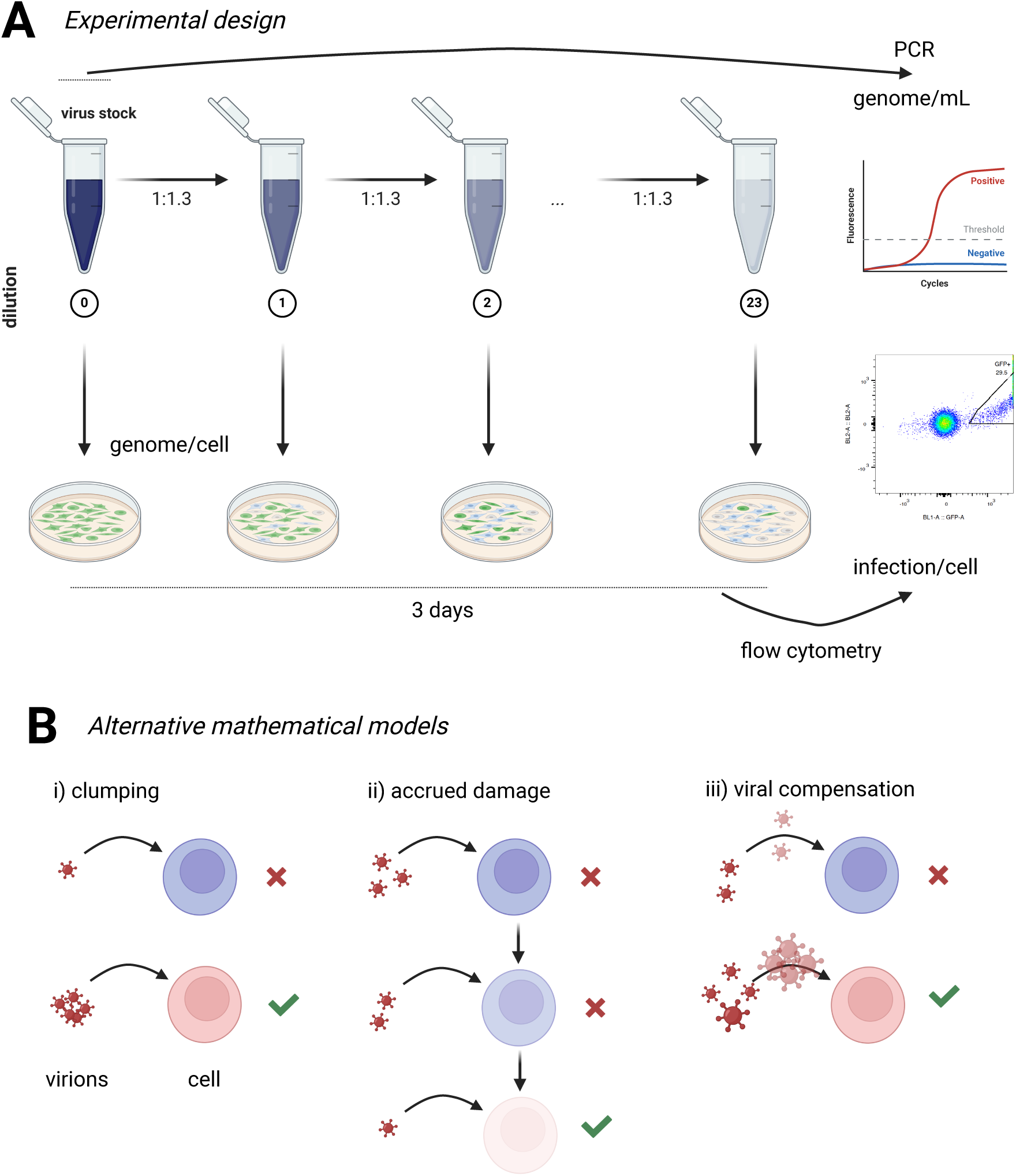
Experimental design and alternative hypotheses for the apparent cooperativity between viruses infecting the same cell. **A**: We diluted stock of different HCMV strains, cloned with reporter genes for GFP or mCherry, up to 23 times and inoculated these dilutions into culture of fibroblasts or epithelial cells. Concentration of virions was determined for each stock using qPCR and the concentration at each dilution was then calculated using the dilution factor in each experiment. We then used flow cytometry to quantify the frequency of target cells expressing GFP or mCherry at 3 days post infection. Kinetics of expression of GFP or mCherry in infected cells over time or at different viral stock dilutions are shown in **Supplemental Figures S2 and S3**. **B**: Three alternative mathematical models aimed to explain apparent cooperativity of HCMV. i) Viral clumping model in which infection of cells mostly occurs when a cell is exposed to a viral “clump” consisting of several, independently acting viral particles with the clump size following a distribution. In the model, virions in a clump attempt to infect the cell independently. ii) Accrued damage model in which exposure of a given target cell by several virions (chosen from Poisson distribution for a given viral concentration) reduces the ability of the cell to resist viral infection (i.e., reduces cell resistance parameter indicated by the level of cell transparency in the cartoon). iii) Viral compensation model in which exposure of a cell to several virions (chosen from a Poisson distribution) with different infectivities increases infectivity of all virions to its highest value in the group of virions.

#### Dynamic Light Scattering (DLS)

Virus stock particle size distributions were measured using a Zetasizer Nano ZS Zen3600 (Malvern Instruments Ltd., Malvern, UK) set at 37^0^C. Virus stocks were diluted 1:3 with DMEM/F-12 without Phenol Red (Gibco, Life Technologies Corp., N.Y., USA), pipetted up and down 10 times to mix, and incubated at 37^0^C for 1 hour. Samples were not filtered, to replicate typical inoculation conditions. Three replicate dilutions were prepared for each stock and analyzed with Zetasizer software (v7.10) using the general size model and viscosity preset for DMEM without phenol.

## Mathematical Models

### Analytical models

To investigate how the probability of cell infection depends on the number of virions/genomes per cell we used a generic “power-law” model similar to our previous work [29, 30]:

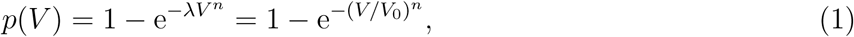

where *V* is number of genomes per cell in the well, and 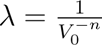 and *n* are model parameters. Parameter *n* denotes a type of interactions between different virions infecting individual cells. Specifically, *n <* 1 implies competition and *n >* 1 implies cooperativity between individual virions for infection, while *n* = 1 indicates that infection of cells proceeds independently (“null” or “single-hit” model, [9, 10, 29, 30]). Infectivity of a single genome is then given as *p*(1) = 1 − e*^−λ^* or *λ* = − ln(1 − *p*(1)). The power-law model is similar to threshold models developed to quantify probability of infection of an organism exposed to different number of infectious particles [31, 32]; however, the threshold models typically display an integer slope at low infection doses that is not consistent with our data.

While the basic “power-law” model (**eqn. (1)**) allows for a saturation in the frequency of infected cells at high genome/cell concentrations it only allows such saturation at one (i.e., lim*_V →∞_ p*(*V*) = 1 in **eqn. (1)**). Furthermore, this model does not describe saturation in the the infection frequency at low genome/cell concentrations. In some of our experiments we observed both: saturation at low genomes/cell (most likely due to few false positive events in the flow cytometer) and saturation below one at high genomes/cell (mechanism is unclear). Therefore, we propose an alternative, 4 parameter model that takes these two limits into account:

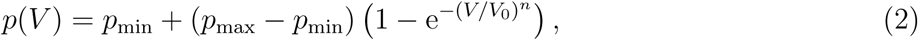

where *p*_min_ and *p*_max_ are the minimal and maximal frequencies of infection observed in a given experiment.

If the relationship between viral inoculum (genomes/cell) and frequency of infection is known (i.e., estimated from the data), the relationship can be used to predict how the infectious titer depends on the inoculum. Infectious titer (in infection units (**IUs**) per mL of culture) is typically found by infecting target cells (given in concentration *C* as cells/mL) with a given concentration of the viral suspension *G* (given, for example, as viral genomes per mL). If the probability of a cell to become infected when exposed to *V* virions (*V* = *G/C* or *G* = *V C*) is given by **eqn. (1)**, infectious titer is then

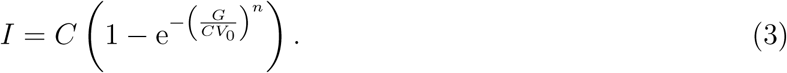

For a given infectious titer *I*, multiplicity of infection (**MOI**) is then calculated as the ratio of the titer to the cell concentration used for infection (*C*_2_):

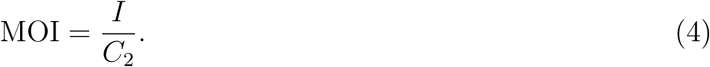

Note that MOI is often calculated for a different target cell cell concentration in experiments than that which was used to titrate the virus stock (i.e., *C* /= *C*_2_) resulting in different numbers of genomes/cell in different experiments.

### Agent-based simulations

To simulate how individual virions infect cells we developed two alternative algorithms: cell-centric and virus-centric. In the cell-centric simulations we focus on individual cells and track if they are infected or not, and in virus-centric simulations we focus on individual virions as to whether it infects a cell or not. Most of our simulations are done using cell-centric approach because ultimately we focus on the probability of a cell to be infected, irrespectively by which virion. We create cell and virion populations by creating cell and virion objects in python or Mathematica. In most simulations, we typically consider 170,000 cells and 16,000-16,000,000 virions per well (these values did vary between simulations to reflect differences in experiments). Each cell and virion is given a randomly-generated “resistance” and “infectivity” levels, respectively, which are taken from a distribution (typically normal but we also investigated other, long-tailed distributions such as log-normal). Most simulations were constructed with Python (ver. 3.7.2) where we defined Cell and Virion classes as follows (see also **Table 1**):

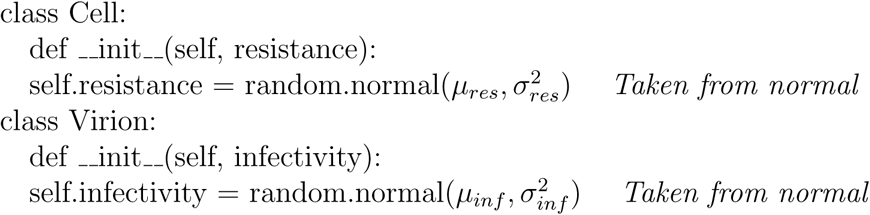

**Table 1:**
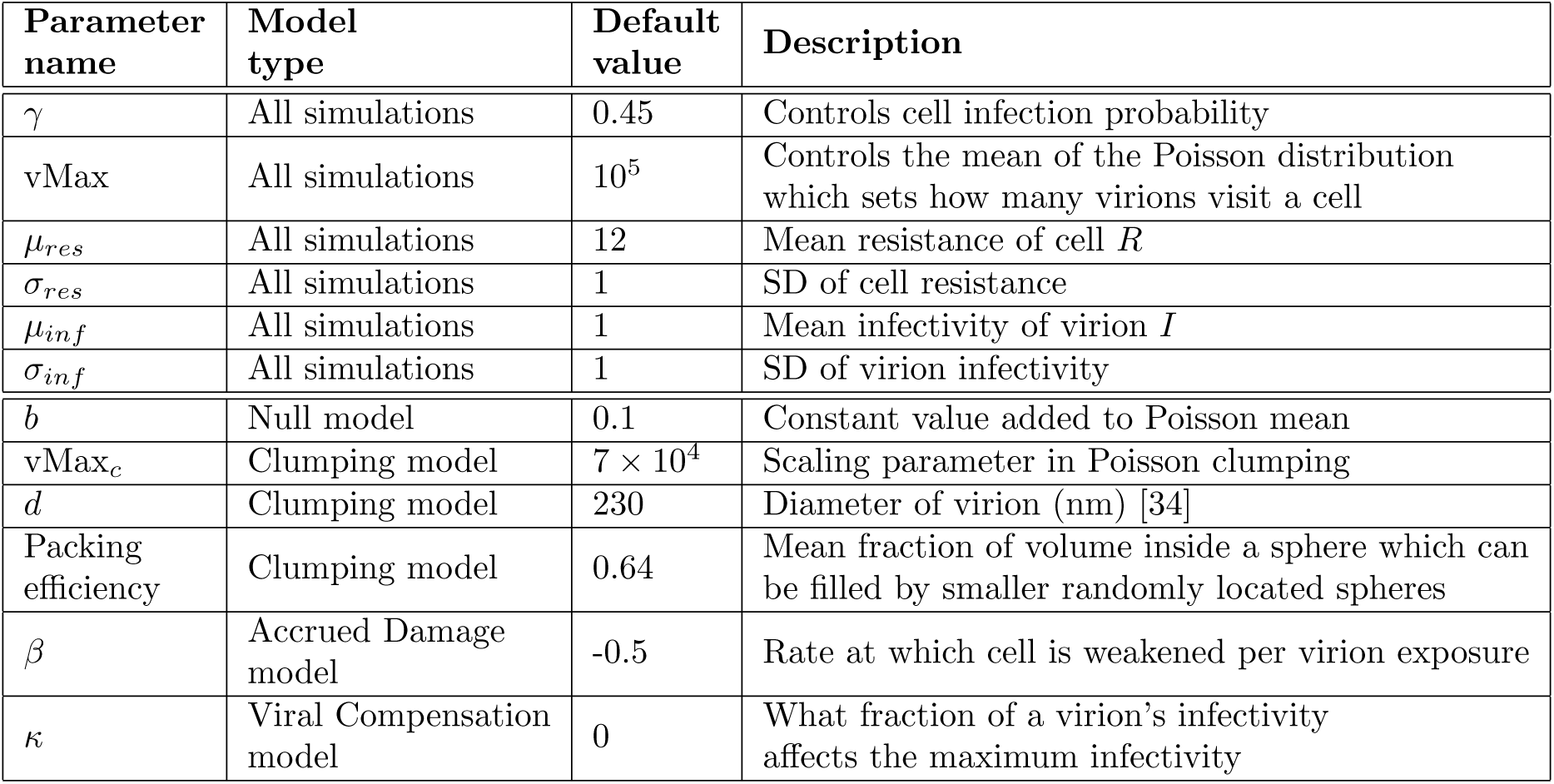
Parameters used in simulations. “Default value” indicates the value of the parameter unless otherwise specified in specific simulations.

| Parameter name | Model type | Default value | Description |
| --- | --- | --- | --- |
| $\gamma$ | All simulations | 0.45 | Controls cell infection probability |
| vMax | All simulations | $10^5$ | Controls the mean of the Poisson distribution which sets how many virions visit a cell |
| $\mu_{res}$ | All simulations | 12 | Mean resistance of cell $R$ |
| $\sigma_{res}$ | All simulations | 1 | SD of cell resistance |
| $\mu_{inf}$ | All simulations | 1 | Mean infectivity of virion $I$ |
| $\sigma_{inf}$ | All simulations | 1 | SD of virion infectivity |
| $b$ | Null model | 0.1 | Constant value added to Poisson mean |
| vMax<sub>c</sub> | Clumping model | $7 \times 10^4$ | Scaling parameter in Poisson clumping |
| $d$ | Clumping model | 230 | Diameter of virion (nm) [34] |
| Packing efficiency | Clumping model | 0.64 | Mean fraction of volume inside a sphere which can be filled by smaller randomly located spheres |
| $\beta$ | Accrued Damage model | -0.5 | Rate at which cell is weakened per virion exposure |
| $\kappa$ | Viral Compensation model | 0 | What fraction of a virion's infectivity affects the maximum infectivity |

In a cell-centric simulation, when a cell is “visited” by a virion, we determine if the cell becomes infected or not using probability function dependent on the difference between values for virion infectivity *I* and cell resistance *R* (**Algorithm** 1). Given the number of virions (or genomes) per cell in a given well, a given cell can be exposed to (visited by) multiple virions. To account for the number of virions a given cell is exposed to, we generated an extra loop and determined the number of virions that would visit the cell from a Poisson distribution whose mean would be a function *f* of the genomes/well. The functions *f* also involves a “vMax” term, a large positive integer, which divides the GENOMES/WELL quantity to make it smaller, so that simulations can be run on a personal computer (**Algorithm** 2). In simulations, every cell is attempted to be infected only once, and virions that had infected a given cell are removed from the virion collection.

#### Algorithm 1

**Cell-centric simulation sketch.** In this scheme, we determine a probability of a cell to become infected when exposed to one virion. When the two interact, their intrinsic infectivity and resistance values are compared, and if the infectivity is greater than the resistance, the cell is infected. Otherwise, a random number between 0 and 1 is generated and compared to the value 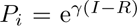 where *I* is the virion infectivity and *R* is cell resistance. If *P_i_* is greater than the random number, then the cell becomes infected, otherwise, cell remains uninfected. In simulations, *γ* = 0.45 unless noted otherwise (**Table 1**).

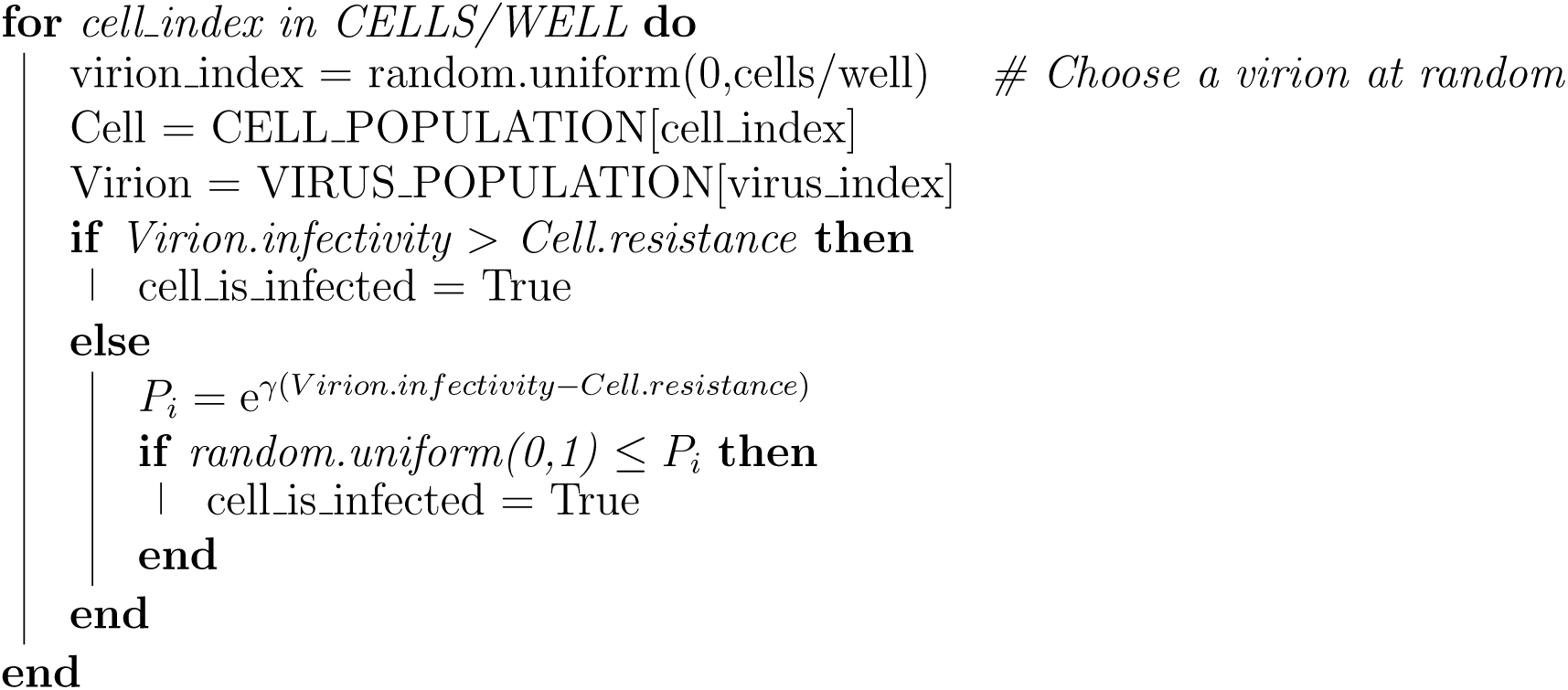

#### Algorithm 2

**Pseudocode to simulate exposure of a cell to multiple viruses.** This code shows a loop implementing the Poisson distribution to determine the number of virions that will attempt to infect a cell. This loop is used inside the cell-centric simulation so that instead of each cell being visited by just one virion, every cell is visited by num virions virions.

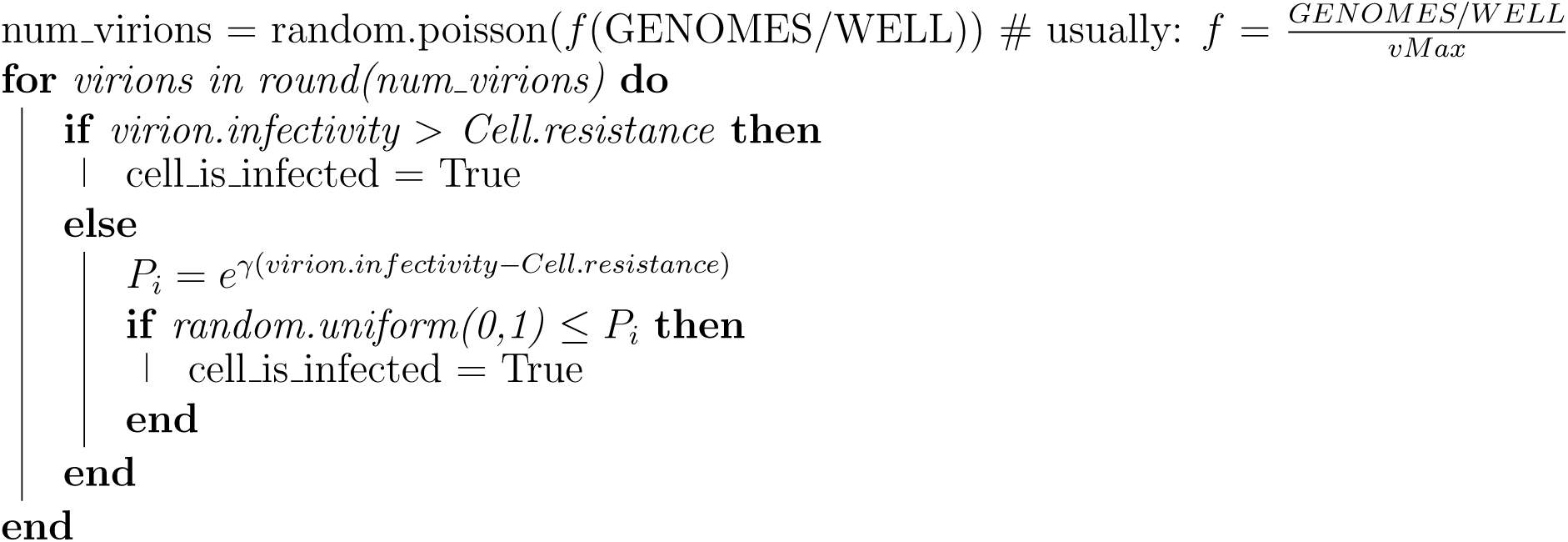

### Clumping hypothesis

In the basic model the number of virions a given cell is exposed to follows a Poisson distribution. However, it is well recognized that as virions are produced by infected cells, they may form clumps/aggregates; the number of virions per clump may deviate from, for example, the Poisson distribution [33]. In the absence of information of what the clump size distribution is, we simulated viral clumps in two alternative approaches. In our python-based simulations, we use SciPy’s minimize function to approximate the ratio of the number of clumps of a given size to the total number of clumps to the Poisson probability density function (**PDF**) value of that clump size. The minimizer has the constraint that the total number of virions per well, i.e. the sum of the number of clumps times clump size, must be equal to the amount of genomes in the well. After obtaining the number of clumps of each size, each virion is assigned a numerical “clump ID”, and if two or more virions share the same clump ID, then they are part of the same clump. The number of clumps of various sizes is determined by Poisson distribution (*P* (*k*)) with estimated mean *λ_c_* where *k* is the number of virions per clump. A clump size of 1 represents a single virion. However, because Poisson distributions have domain 0, 1, 2, 3*, …*, we took 0 to represent a clump size of 1, 1 to represent clump size 2, and so on (**Algorithm** 3).

#### Algorithm 3

**Pseudocode to model viral clumps.** In this model, clump distributions and clump IDs are created according to a Poisson distribution, *P* (*i*). First, the maximum clump size is chosen by locating the integer where the CDF of the Poisson distribution is greater than 0.99. Next, we choose the number of clumps of size 1, 2*, . . .,* max such that the ratio of the number of clumps of size *i* divided by the total number of clumps in the sample is as close to *P* (*i*) as possible. To do this, we use minimize the error between *P* (*i*) and 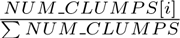 using the negative log likelihood error function NegLogLike in python. After the best values of *NUM CLUMPS*[*i*] are obtained, we assign every clump a unique integer ID which applies to every virion inside that clump.

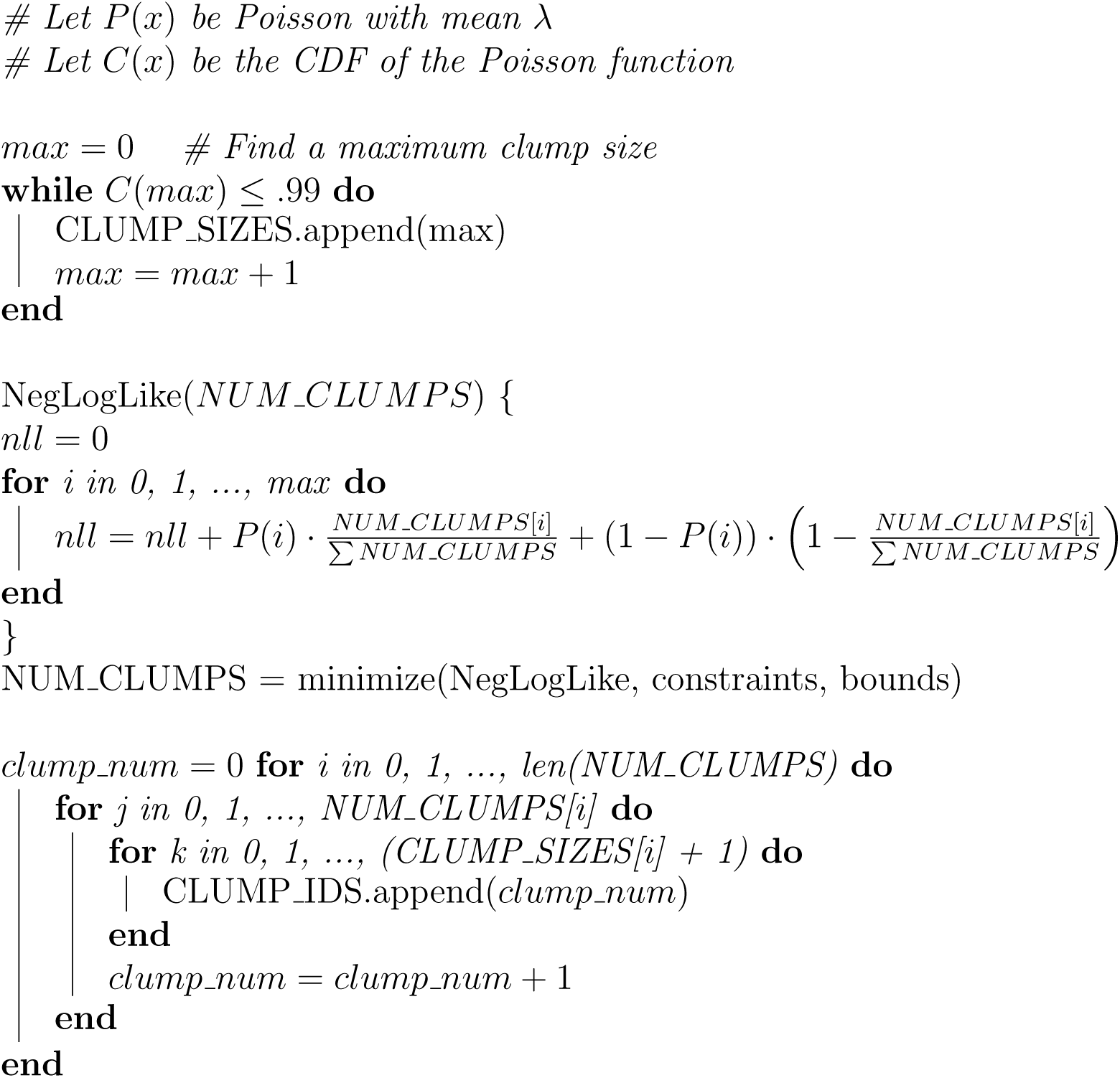

In our alternative, Mathematica-based simulations, we used one-inflated geometric distribution to simulate the number of virions in different clumps, we varied the fraction of clumps that would have one genome and allowed other clumps to have larger mean size (e.g., 50 virions/clump). Specifically, we assumed that a fraction *f*_1_ of clumps have exactly 1 virion and 1 − *f*_1_ clumps follow geometric distribution *P* (*k*) = *p*(1 − *p*)*^k^* with *k* = 1, 2*, . . .* where *p* determines skewness of the distribution. We then assigned every virions in the library of 10^5^ virions to each clump, and in simulations, when *v* virions attempt to infect cells (chosen from Poisson distribution for a given dilution of the stock), all virions belonging to the clump IDs of the infecting virions also attempt to infect the cell.

In the clumping model, we performed two sets of simulations – in one set if one virion from a clump is successful at infecting a cell, the clump (along with all virions associated with that clump) is removed from the well. In the alternative simulations we allowed the clump to remain, with virions in that clump being able to infect other cells.

To complement our theoretical assumption that clump size distribution follows Poisson or geo-metric distribution, we also used experimentally measured distribution of clump sizes for a given viral stock dilution (**Supplemental Figure S10**). For infection of a cell, we sampled the clump size from experimental distribution (**Supplemental Figure S11**) and then calculated the number of virions in the clump using

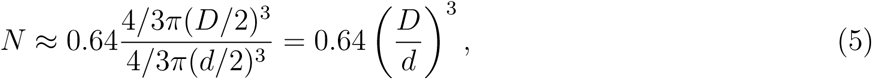

where *D* is the clump diameter, *d* = 230 nm is the diameter of HCMV virion [34, 35], and 0.64 is the packing efficiency of randomly distributed spheres [36, 37]. The number of virions per clump scales nonlinearly with the clump size, i.e., a clump of size *D* = 2*d* = 460 nm would have 5 virions, and clump of size *D* = 3*d* = 690 nm would have 17 virions.

### Accrued damage hypothesis

Exposure of a cell to a virion may trigger generation of antiviral response that may reduce the overall resources the cell has. We hypothesize that for each virion that attempts to infect a cell but fails to do so, the level of cell resistance *R* decreases by a value *β*. This effectively increases infection chances of the next virion in the collection of virions attempting to infect a given cell.

### Viral compensation hypothesis

In this scheme, a collection of virions will attempt to infect a cell. We increase infectivity of all infecting virions in the collection to a value proportional to the maximal infectivity of virions in the collection max(*I_j_*) and the current infectivity of the virion *I_i_* as

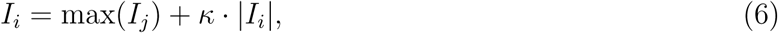

and in most simulations we allowed *κ* = 0 (i.e., infectivity of each virion in the collection of all virions attempting to infect a cell increases to a maximum value in the collection, **Table 1**).

### Virus-centric simulations

We also simulated virus infection using a virus-centric approach, where each virion in the initial pool determined by genomes/well in the data, gets one chance to infect a cell. These virus-centric simulations gave the same prediction on how the probability of cell infection scales with the number of virions per cell for the null model and thus are not presented here.

## Statistics

To estimate parameters of the models relating the cell infection probability and the number of virions in culture we use a likelihood approach [29, 30]. Specifically, the negative log-likelihood of the model (e.g., **eqn. (1)**) is

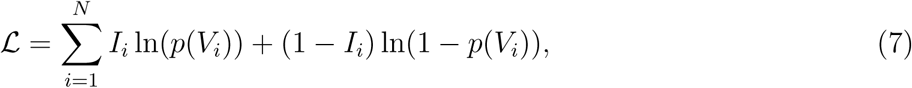

where *V_i_* and *I_i_* are the number of genomes (or virions) per cell and the proportion of infected cells in the well, respectively, and *i* denotes sample. Because most data did not exhibit a linear relationship between virion concentration and infection probability we fitted the models to subsets of data best matching a linear relationship on a log-log scale. Note that linear relationship on log-log scale may still be nonlinear (on linear-linear scale) when *n* /= 1. We also fitted an extended power-law model (**eqn. (2)**) that accounts for saturation in infection probability to the datasets that include all data points. We used the same approach to estimate degree of apparent cooperativity *n* for the “synthetic” data that we generated in simulations.

## Results

### Infection of target cells by fluorescently-tagged HCMV strains often deviates from a single hit model

In our previous studies, we showed that the specific infectivity (**SI**) values of several strains of HCMV were dependent both on the choice of target cell type and on the cell type used to produce the virus stock [4]. We also noted that even when the same producer cell type and target cell type were used, titer and, consequently, SI values could differ between assays performed with different inoculation concentrations (e.g., see Graphical abstract). This concentration-dependent effect suggested that virions may be either cooperating or interfering with each other while infecting a target cell. To examine whether these differences were caused by cooperation between virions (i.e., during coinfection of target cells by multiple virions), we performed small-step titration assays to produce high-resolution dose-response curves for HCMV strains TB, ME, and TR, on two target cell types often used in research: human foreskin fibroblasts (fibroblasts) and ARPE-19 retinal pigment epithelial cells (epithelial cells or **ECs**). Our three virus strains express either GFP or mCherry proteins in place of the US11 gene and the rate of infection was determined by counting either GFP+ or mCherry+ cells by flow cytometry at three days post infection (dpi) [23]. We chose to quantify cell infection frequency at 3 dpi because HCMV progeny virions are typically released around day 3 [38–40] and even though expression levels of GFP or mCherry in cells infected by progeny virus would typically be very low at 1 dpi, robustly detecting infected cells at this time may not be fully feasible in our experimental system (**Supplemental Figures S2 and S3**). Furthermore, we noted that cell infection frequency did not strongly depend on the genome/cell concentration at very low genomes/cell (e.g. TB-GFP on fibroblast, **Figure 2A**) further suggesting that such low frequency events may contain false positives (**Supplemental Figures S2 and S3**). Yet, for many strain-cell combinations, cell infection frequencies of ≥ 10^−4^ were robustly detected (**Figure 2**).

**Figure 2:**
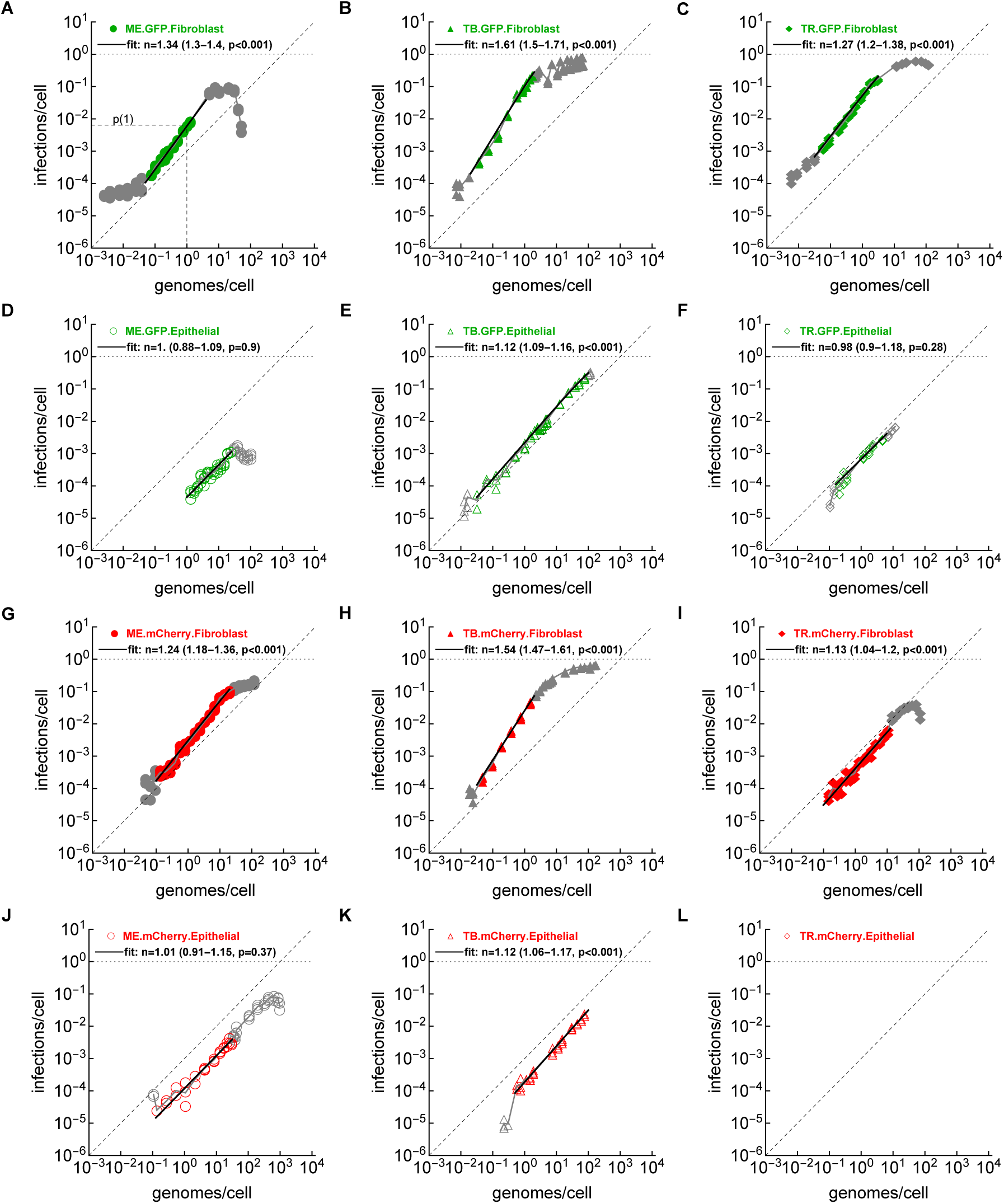
Apparent cooperativity of human CMV strains at infecting different target cells. We performed experiments by infecting either fibroblasts (HFF) or epithelial cells (ARPE-19) with three different strains of HCMV (ME, TB, TR) expressing two different reporter genes (GFP or mCherry, see **Figure 1A**). We then fitted a “power-law” model (**eqn. (1)**) to subsets of these data (shown in color) and calculated parameter *n*, indicating a degree of apparent cooperativity of the virions at infecting cells (*n* = 1 – no cooperativity, *n >* 1 – apparent cooperativity, see Materials and Methods for more detail). For each estimate we also show *p* value from the likelihood ratio test when comparing the power-law model fit to the single-hit (*n* = 1) model fit. The solid black lines show the predictions of the mathematical model (**eqn. (1)**) in the range of data that was used in model fitting, the data not used in fitting are shown in gray, and empty panel in **L** denotes an experiment that could not be performed due to lack of cell infection (TR.mCherry on ECs). All experiments were performed at least twice. Data are shown as markers and model fits as black lines. Diagonal dashed lines represent the slope of one. In panel A, we show infectivity of a single genome, *p*(1) from **eqn. (1)**. Fits of the extended power-law model (**eqn. (2)**) are shown in **Supplemental Figure S4**.

All strain-cell combinations displayed saturation at high inoculum concentrations, although none achieved a higher than 98% infection rate. This is not unexpected, as we did not synchronize cell cycle before inoculation and some cells may not be fully susceptible to infection. Also, exposure to a large number of virus particles may render some cells highly resistant to infection [12]. Interestingly, we noted a decline in the infection frequency at higher concentrations for ME and TR strains but not for TB (**Figure 2**); such a decline in the infection frequency is inconsistent with a single hit model.

To more rigorously characterize deviation of the infection rate from a single hit (null) model, we fitted a “power-law” model that allows for a nonlinear change in the infection rate with increasing genomes/cell to the data (see Materials and methods for detail). In this analysis, we sub-selected data that fall on a line on a log-log scale and estimated parameter *n* characterizing how cell infection rate scales with the larger virions/cell and parameter *p*(1) representing infectivity of a single genome (**Figure 2A**). The range for which the log of infections/cell scaled linearly with log of genomes/cell varied by specific strain-cell type combinations was only 1 order of magnitude for ME-GFP on epithelial cells (**Figure 2D**) and 3 orders for TB-GFP on epithelial cells (**Figure 2E**). Importantly, for most HCMV strain-target cell combinations we estimated *n >* 1 (**Figure 2 and Supplemental Table S2**). With *n >* 1 increase in virion concentration (i.e., higher genomes/cell values) results in a higher than linear increase in the probability of a cell to be infected (**eqn. (1)**) indicating cooperation between virions at infecting cells. We call this phenomenon “apparent cooperativity”. Fitting a more complex model that allows for limit of detection of infected cells at low genomes/cell and saturation in the infection rate at lower than one values at high genomes/cell also resulted in *n >* 1 for most strain-cell combinations although there were clear biases in the model fit of some data (**Supplemental Figure S4**). We also found *n >* 1 when fitting the power-law model (**eqn. (1)**) to subsets of data defined by a sliding window further supporting existence of apparent cooperativity for a wide range of genomes/cell values (**Supplemental Figure S5**). Interestingly, we also found evidence of apparent cooperativity after fitting the power-law model to data available on infection of target cells by HIV and vaccinia virus, but not on formation of viral plaques on leaves inoculated with tobacco mosaic virus (**Supplemental Figures S6 and S7**).

While the degree of apparent cooperativity *n* and infectivity of a single genome *p*(1) was de-pendent on the specific strain-cell type combinations, some patterns were evident. In particular, all strains had a higher degree of cooperativity and higher genome infectivity on fibroblasts than those on epithelial cells (**Figure 3A&D**). HCMV TB was the strain with a highest genome infectivity *p*(1) (**Figure 3F**) and demonstrated highest degree of cooperativity (**Figure 3C**); we detected the highest degree of apparent cooperativity for TB-GFP on fibroblasts (*n* = 1.61, **Figure 2B**). Importantly, the type of the reporter gene had no influence on either *n* or *p*(1) suggesting that a nonlinear relationship between virion concentration and cell infection rate was not due to different virus modifications (**Figure 3B&E**). Interestingly, when pooling all estimates together there was a strong positive correlation between virion infectivity *p*(1) and degree of apparent cooperativity *n* (**Figure 3G**). Taken together, our careful titrating of stocks of three different strains of HCMV revealed a highly nonlinear relation-ship between virion concentration and the cell infection probability that deviates from predictions of a commonly assumed single hit model.

**Figure 3:**
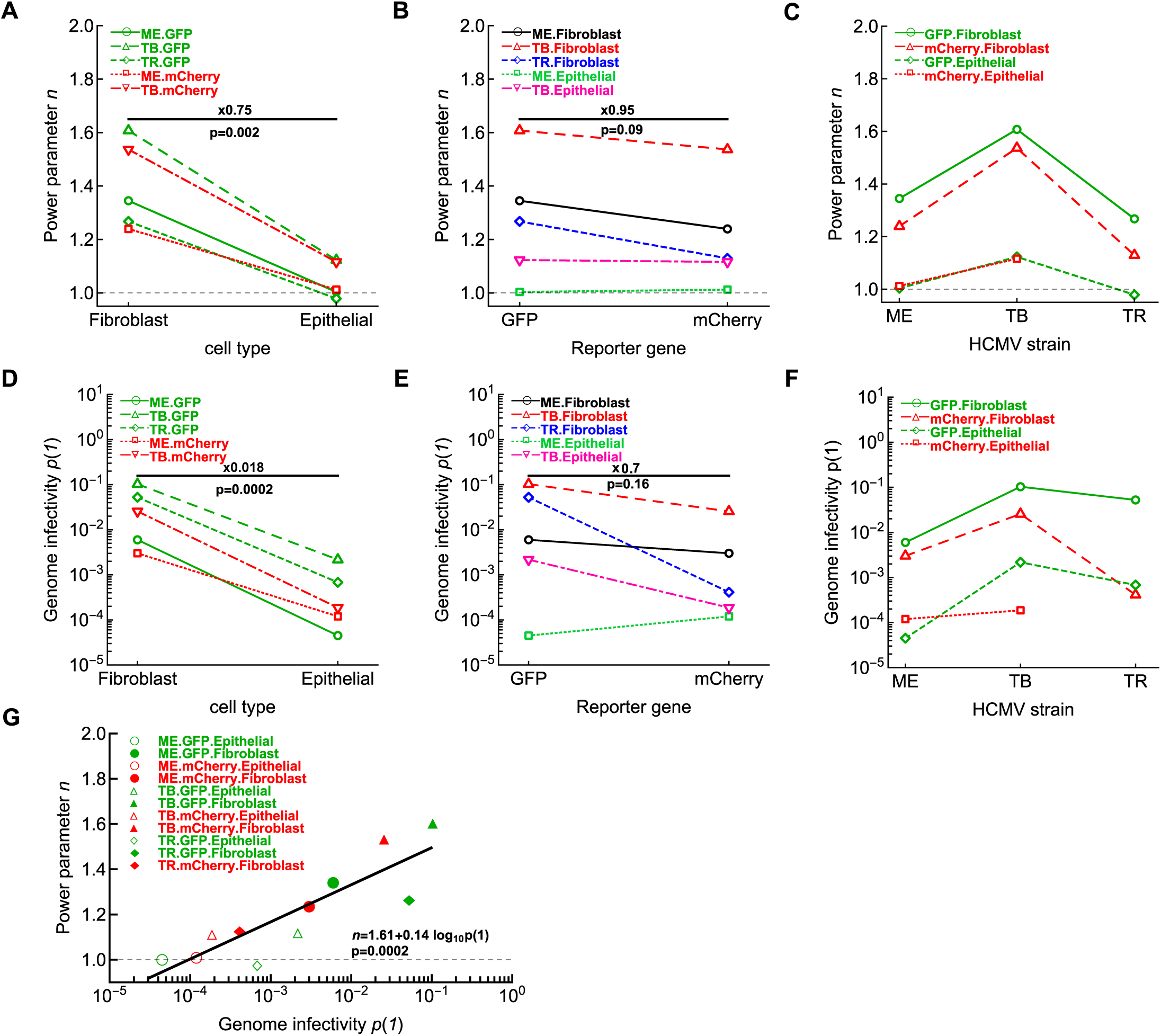
All HCMV strains show reduced infectivity and reduced apparent cooperativity when infecting epithelial cells. We plot the relationship between estimated power parameter *n* (**A**-**C**) or virion infectivity (infectivity of a single genome, *p*(1), **D**-**F**), and other viral and cell characteristics such as target cell type (**A**&**D**), reporter gene (**B**&**E**), and HCMV strain (**C**&**F**). Parameter *n* indicates cooperativity in virus infection of cells (estimates are given in individual panels in **Figure 2**). Symbol ‘x’ indicates fold difference, and listed *p* values are from paired t-test. In panel **G** we show the relationship between estimated virion infectivity *p*(1) and apparent cooperativity *n* estimated for different strain/cell combinations; we also show the linear fit and *p* value from the test of no relationship.

### Random infection model cannot explain apparent cooperativity of HCMV strains

Finding that a probability of cell infection increases faster than linearly with virion concentration was unexpected and to explore potential mechanisms of such apparent viral cooperativity we created a computational model in which we track infection of cells by virions (see **Algorithms 1–2**). We created an array for cells, each with different level of resistance, and an array of virions, each with different degree of infectivity [41]. In this model we attempted to mimic experiments by using similar (but scaled) numbers of target cells per well and by varying the number of virions present in a well. We then simulated dilution of viral stock resulting in different numbers of virions/well (and thus, virions/cell) and by using Poisson distribution calculated the number of virions each cell in the population would be exposed to (**Algorithm** 2). Assuming that each virion attempts to infect a cell independently of other virions and considering that infection event is a random process dependent on actual values of cell’s resistance and virion’s infectivity (**Table 1**) we found that this random infection/null model predicts linear increase in the cell infection probability with increasing virions/cell (**Figure 4A**), as predicted by a single hit model.

**Figure 4:**
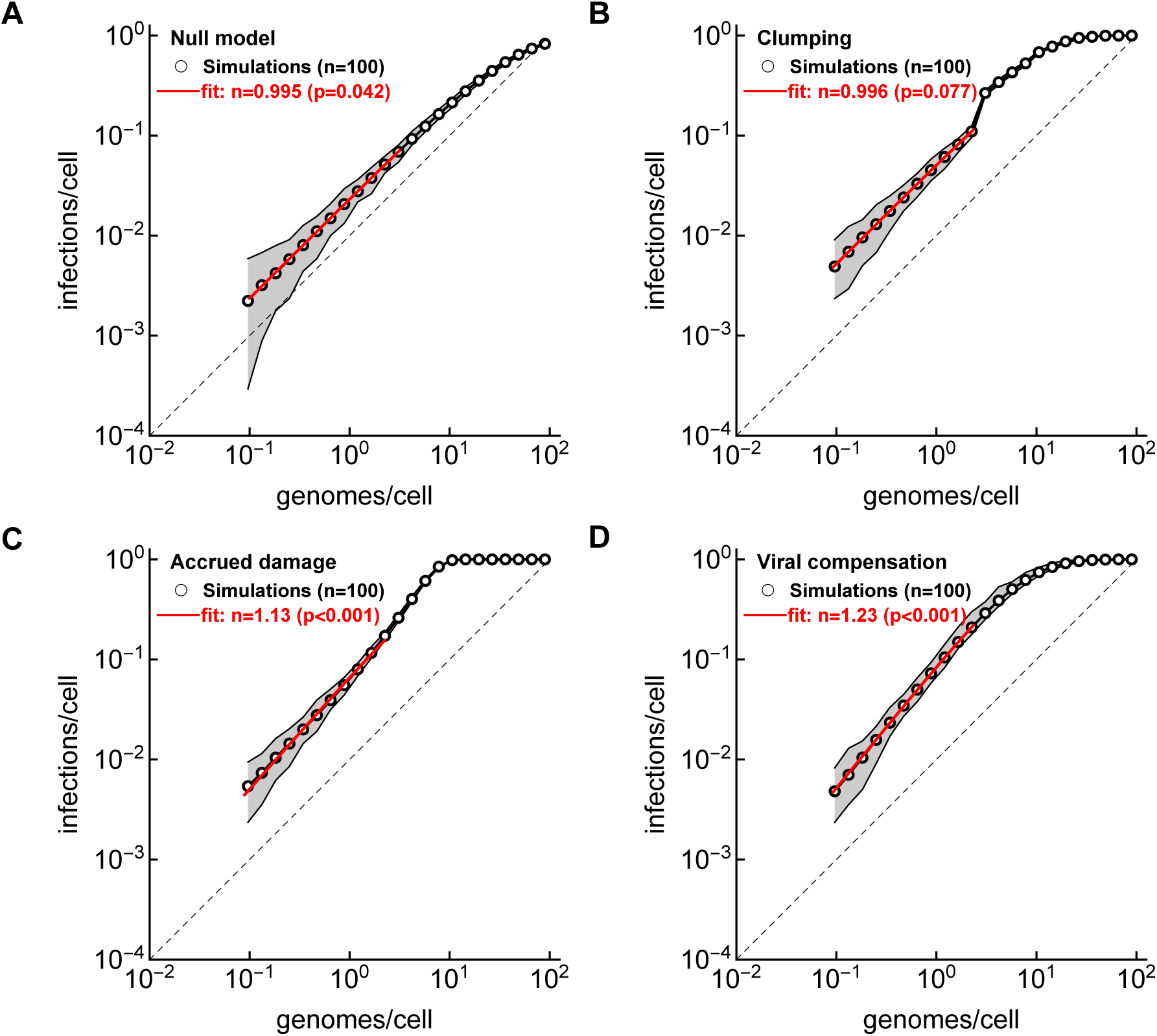
Heterogeneity in virion infectivity and/or susceptibility to infection of target cells and aggregation of virions does not result in apparent cooperativity in simulations. We developed simulations in which individual virions vary in their infectivity and individual cells vary in their resistance (or susceptibility) to infection (see Materials and methods for detail). We then sampled individual virions and cells randomly assuming different ratio of virions (genomes) per target cell and calculated how frequency of infected cells changes with increasing number of viral genomes per cell for a null model (**A**), clumping model in which virions aggregate (**B** and **Figure 1Bi**), accrued damage model, in which individual virions attaching to a cell reduces its resistance (**C** and **Figure 1Bii**), and viral compensation model in which virions attaching to a cell acquire infectivity of the most infectious virion (**D** and **Figure 1Biii**, see Materials and methods for more detail). We simulated infection of target cells by virions according to these models and searched for parameter combinations that qualitatively matched experimental data for HCMV-TB-GFP infection of fibroblasts (with *n* = 1.60, **Figure 2B**). We fitted the power-law model (**eqn. (1)**) to the results of simulations using likelihood approach (**eqn. (7)**) and estimated the degree of apparent viral cooperativity (*n*) for linear parts of the data at low genome/cell values (shown by red lines). Averages are shown by markers and gray areas indicate the range of values in simulations. Parameters used in simulations are given in **Table 1**. Note that to model viral clumps we assumed that clump size followed Poisson distribution; other details are given in Material and methods section.

We have tried multiple combinations of parameters in our simulations and also developed simulations from “virus-centric” perspective; none of the simulations resulted in apparent cooperativity with *n >* 1. Because it is possible that we may have missed just the right set of parameters that would give us indication of apparent cooperativity, we turned to analytical derivations. Let us assume that infection of target cells follows a mass action law, *p*(*V*) = 1 − e*^−λV^* (i.e., with *n* = 1 in **eqn. (1)**); here *λ* indicates susceptibility in the target cell to the infection. Let us also assume that target cells vary in their susceptibility to infection according to a distribution *f* (*λ*). Then the probability of a cell to be infected when exposed to *V* virions/genomes is

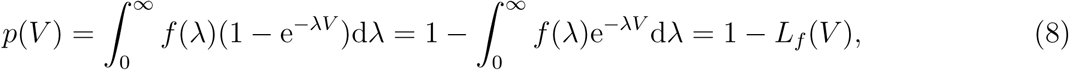

where *L_f_* (*V*) is the Laplace transform for *f*. Now we can use Taylor’s expansion for *L_f_* (*V*)

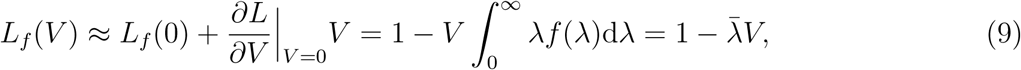

where 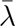 is the average of the distribution *f* (*λ*). Then the infection probability from **eqn. (8)** becomes genome/cell concentrations, the probability of infection scales linearly with genome/cell. This result is independent whether virions vary in their infectivity.

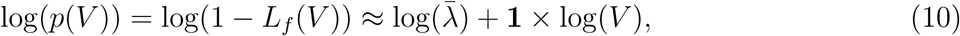

### Three alternative hypotheses to explain apparent cooperativity

We investigated what alternative mechanisms may result in deviation of cell infection from that predicted by the single-hit model. One hypothesis was that individual virions may not simply infect a cell independently but form aggregates (or clumps) and such clumping may increase chances to infect a cell (**Figure 1A** and [16]). We therefore extended our basic model to allow virions to clump, and in this first “clumping model” we assumed that clump size distribution follows a Poisson distribution with the mean that was scaled (with a parameter vMax*_c_*) to the calculated number of genomes/well (**Table 1**). We then used Poisson distribution to sample clumps and allowed virions in the clump to independently attempt to infect a cell. In contrast with the null model, the Poisson clumping model predicted nonlinear change in the frequency of infected cells with a large increase in infection frequency at about 2 genomes/cell (**Figure 4B**). However, at lower genomes/cell values, the model predicted linear relationship between infection frequency and virion concentration, similar to the null model. Thus, virions clumping, at least at these tested model parameters, did not result in apparent cooperativity at small genomes/cell values. We note that in these simulations we did not remove clumps from the well if one of the clump’s virions succeeds at infecting a cell; extending the simulations to allow for clump removal upon successful infection did not change the results of the clumping model (**Supplemental Figure S8**).

We next explored alternative hypotheses that include some type of “memory” of sequence of interactions between a target cell and individual virions. In particular, we found inspiration in the remarkable experiments by Stiefel *et al.* [42] in which the authors discovered that sequentially depositing two or more vaccinia virions on individual target cells resulted in the disproportionally higher probability of a cell to become infected; we estimated a high degree of apparent cooperativity (*n* = 2.24) in these experiments (**Supplemental Figure S6B**). One explanation of these results became our “accrued damage” hypothesis in which each interaction between a cell and a virion that does not result in infection does lead to increased susceptibility of the cell upon encounter with another virion (**Figure 1B**). This may arise due to depletion of resources a cell must use to prevent productive infection upon interaction with a virion. We added this mechanism in our simulations by allowing resistance of the cell *R* to be reduced by *β* at each virion encounter until either the cell becomes infected or no more virions attempt to infect the cell. Even for a relatively small *β*, model simulations did result in apparent cooperativity at low genomes/cell values and increased even faster at intermediate genomes/cell (**Figure 4C**). Interestingly, the overall shape of the relationship between the frequency of cellular infection and viral concentration (genomes/cell) visually matched that observed experimentally for TB-GFP strain of HCMV infecting fibroblasts (**Figure 2B**).

Our accrued damage model suggests that interactions between virions and cells result in changes in the cell that lead to its increased susceptibility to infection. However, an alternative hypothesis is that when a cell is exposed to several virions, virions may be augmenting their infectivities by complementing different functions required for infection (**Figure 1B**). This “viral compensation” hypothesis has been previously suggested following observations of increased infectivity of viral aggregates [16, 33, 43]. We therefore investigated whether virions may “cooperate” at infecting target cells even if they do not aggregate but when the number of virions attempting to infect an individual cell is described by the Poisson distribution. In our version of viral compensation model we gather information on the collection of virions attempting to infect a given cell and re-assign infectivity to each virion depending on the maximal virion infectivity present in the collection (see **eqn. (6)** and Materials and methods for more detail). In simulations where all virions (generated by a Poisson distribution) attempting to infect a cell increase their infectivity to the maximal value in the collection, the frequency of cellular infection increased faster than linearly with genomes/cell, i.e., results in apparent cooperativity (**Figure 4D**). Thus, our results so far suggest that apparent cooperativity may result in the non-Markovian process of cell infection either by individual virions (accrued damage model) or because of compensation of ability to cause productive infection by virions attempting to infect the same cell (viral compensation model). Interestingly, we could find sets of parameters that allowed for these two alternative models match approximately experimental data (**Supplemental Figure S9**).

### Direct testing of the viral clumping hypothesis as explanation for apparent cooperativity of HCMV at infecting targets

Intuitively, it makes sense why accrued damage and viral compensation models may result in apparent cooperativity with *n >* 1 at low genomes/cell values: as more virions attempt to infect a cell, the cell loses resistance quicker (accrued damage model) or virions have higher chance of increasing infectivity (viral compensation model) leading to higher probability of cell infection. It was less clear why a viral clumping model did not lead to apparent cooperativity. A number of previous studies documented increased infectivity of viral clumps as compared to individual virions [16]; yet, we were not able to find studies that rigorously tested if viral clumping results in apparent cooperativity at low genomes/cell values.

Presence of viral aggregates is common in viral stocks [16]. For example, by using electron microscopy Galasso & Sharp [33] quantified the number of vaccinia virions present in different clumps/particles; 67% of particles were single virions and the largest particle had 81 virion. The resulting distribution of virions per clump was highly skewed with the average 3.2 virions per clump [33]. HCMV can also form aggregates but may also exist as single particles [34]. To study how dilution of our HCMV stock may influence distribution of viral clumps we utilized dynamic light scattering (**DLS**) to measure the distribution of particle sizes in stocks of all three strains [44–46]. For the stock of TB-GFP strain of HCMV grown on fibroblasts we found that the distribution of particle diameter *D* depends strongly on the stock dilution; at low dilutions (i.e., dilution #1-4) nearly all particles were larger than 100 nm but at higher dilutions (e.g., dilution #19), particles with the average size of 20 nm became dominant (**Figure 5A and Supplemental Figure S10**). The peak at larger sizes likely represents virions and given that an average HCMV virion diameter is *d* = 230 nm [34], some of the smaller clumps represent viral particles without capsid, cellular debris, exosomes, and particles already present in the growth media. By fitting a mixture of two normal distribution to log-transformed clump size data we estimated proportion of the larger clumps (*f*_1_) and their average scaled diameter (*D̅/d*); the model in which the average of the larger distribution remained constant between dilutions but their fraction declined with higher stock dilution described the data reasonably well (**Supplemental Figure S10**).

**Figure 5:**
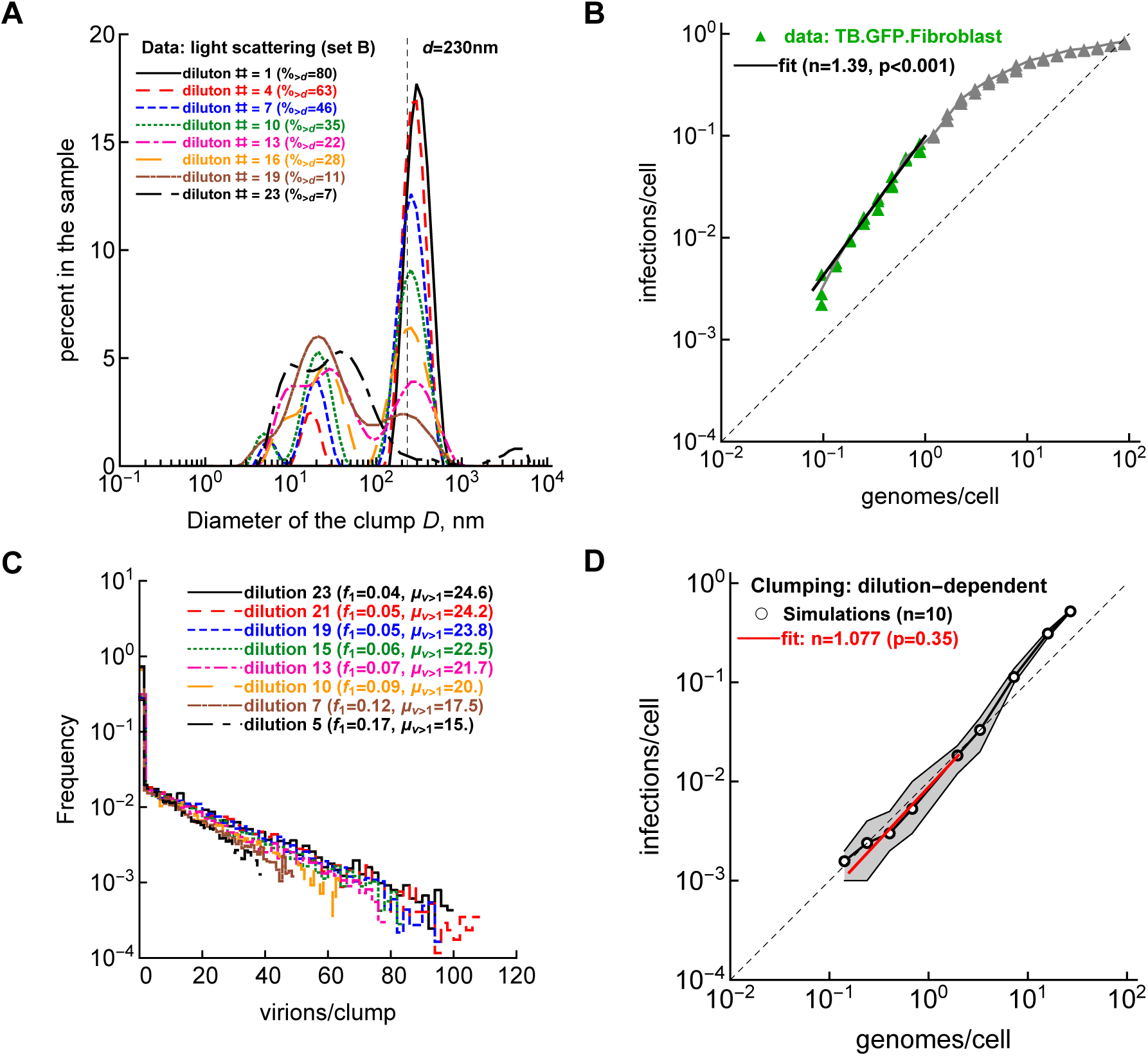
Incorporating experimentally measured distribution of viral aggregates in simulations did not result in apparent cooperativity in the clumping model. **A**: We used dynamic light scattering to measure distribution of sizes of clumps for various dilutions of the initial stock of TB-GFP strain (see also **Supplemental Figure S10**). **B**: We used the same dilutions of TB-GFP from the same cultures to infect fibroblasts and estimated the fraction of infected cells at day 3 after infection. We then fitted the power-law model (**eqn. (1)**) to the linear part of the data and estimated degree of apparently cooperativity *n*. **C-D**: We used experimentally measured distribution of clump sizes and estimated how many virions are located in a given clump of a diameter *D* assuming random packing of spheres with *d* = 230 nm (see **eqn. (5)** and text in Materials and methods). We then simulated infection of cells by sampling virions in clumps (clumping model, **C**) and calculating probability of cell infection at different genome/cell dilutions. We show the results of 10 simulations with parameters listed in **Table 1** except we used 10^3^ target cells. In **D**, we fitted the power-law model (**eqn. (1)** to the data from simulations and estimated the degree of apparent cooperativity *n*. Makers denote average values and gray areas show the range of simulation results. Listed p-values were calculated to estimate deviation of estimated *n* from *n* = 1 using LRT. The size of HCMV virion of *d* = 230 nm is indicated by a dashed vertical line in panel **A**.

We then used the same stock dilutions to infect fibroblasts and estimated the degree of apparent cooperativity; as in previous experiments we found strong deviation of the infection from the null model with a high degree of apparent cooperativity (*n* = 1.39) for genomes per cell changing over an order of magnitude (**Figure 5B**); notably estimated *n* in these experiments was lower than that we observed earlier (**Figure 2B**). To investigate whether experimentally measured clump size distribution may be able to result in apparent cooperativity in simulations, we used a previous result suggesting that a number of smaller spheres that could fit inside of a larger sphere scales as cube of the ratio of diameters of the two spheres (**eqn. (5)** and **Supplemental Figure S11A**). By using fit of the mixture of two normal distributions we estimated how many virions are predicted to be in the well at a given dilution and how the number of virions per clump changes with dilution of the stock (**Supplemental Figure S11B-C**). Interestingly, while the total number of virions declines with higher dilution number, the proportion of clumps with more virions increases with higher dilutions. To allow for skewed distribution of the number of virions per clump we used one-inflated geometric distribution (see Materials and methods for detail and **Figure 5C**). Interestingly, by assuming dilution-dependent clump size distribution we found that the cell infection rate scales nearly linearly with increasing virion concentration **Figure 5D**). However, these simulations also revealed small (but statistically non-significant) evidence of apparent cooperativity at small genomes/cell values (*n* = 1.08) and even faster increase in infection frequency at higher virion concentrations (**Figure 5D**).

We performed many additional simulations with the clumping model by varying the distribution of virions per clump, how clumps and virions are sampled at a given stock dilution, and changing levels of virion infectivity. We found that in some simulations, specifically when the frequency of clumps with many virions was sufficiently large at high dilutions, the clumping model could result in apparent cooperativity with *n >* 1 (e.g., **Supplemental Figure S12**). These results suggest that understanding how dilution of the stock impacts the distribution of clump sizes may influence of how cell infection frequency scales with virion concentration. In contrast, additional simulations with the accrued damage and viral compensation models (with different parameter values and different simulation frameworks) nearly always resulted in apparent cooperativity, suggesting that these two hypotheses are more robust to modeling details. In fact, we could find parameter combinations that allowed for relatively high values of the cooperativity parameter *n*, e.g., in the accrued damage model assuming larger reduction in cell resistance per virion encounter (e.g., *β* = −2) easily led to *n* ∼ 2. Taken together, our results suggest that both accrued damage and viral compensation models, and under some parameters, viral clumping model, may allow for the cell infection rate to increase more rapidly with higher virion concentrations than predicted by the single hit model.

### Calculations of MOI are imprecise for virus-cell combinations with apparent viral cooperativity

Our experiments and computational models show that for some virus-target cell combinations the frequency of cell infection increases faster than linearly with higher genomes/cell values. Even though we were not able to discriminate between alternative hypotheses for such apparent viral cooperativity, such nonlinear scaling may impact calculation of the infectivity of a viral stock and may influence interpretation of experiments involving comparison of different viral strains.

To calculate titer of a given stock, the stock is diluted so the number of infectious units (**IUs**) can be calculated. In case of a linear, single hit model, the estimated titer (IU/mL) scales linearly with the virion concentration in the sample independently of the concentration of target cells used in the experiment (**Figure 6A**). In contrast, in the presence of apparent cooperativity, estimated titer even when measured at the same virion concentration strongly depends on the concentration of target cells (**Figure 6B**) making calculation of stock titer imprecise. This is because as virus and/or target cell concentrations vary, the ratio of virions/cell also varies resulting in different predicted levels of infection.

**Figure 6:**
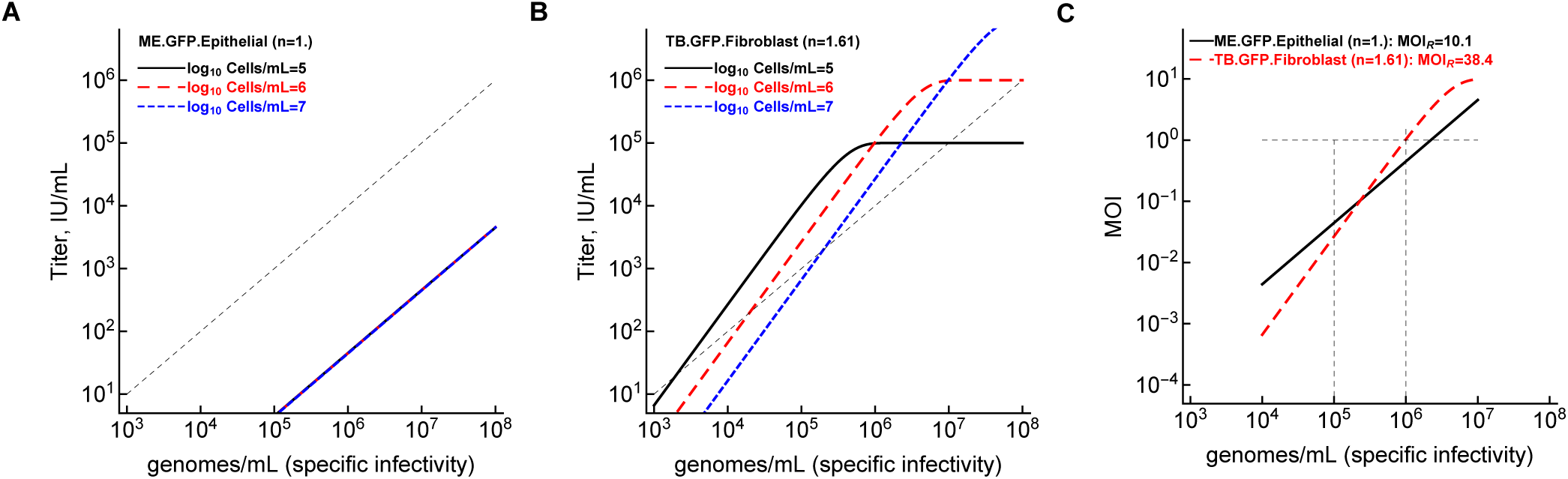
Calculations of MOI are imprecise for virus-cell combinations exhibiting apparent coopera-tivity. **A**: For viruses with *n* = 1, infectious titer scales linearly with changing genomes/cell (ME.GFP.Epithelial). **B**: However, for viruses with *n >* 1 (TB.GFP.Fibroblast), reduction in genome/cell results in a larger reduction in infections/cell, i.e., in much lower than expected infectious titer. **C**: Expected MOI changes nonlinearly for virus-cell combinations with apparent cooperativity (*n >* 1); MOI*_R_* is the ratio of MOIs when specific infectivity is reduced 10 fold (i.e., stock is diluted 10 fold, from 10^6^ to 10^5^ genomes/mL); here target cell concentration is 10^6^ cell/mL. In panels A and B, dashed line denotes 1:1 relationship between genomes/mL and infectious titer (IU/mL). We used parameters of the power-law model fitted to individual datasets (for ME-GFP on ECs and TB-GFP on fibroblats) to make predictions (**Supplemental Table S2**).

If different viral strains vary in their degree of apparent cooperativity (characterized by the parameter *n*) then some comparisons of viral characteristics, e.g., specific infectivity or replicative capacity, may be difficult to interpret. For example, it is common to study replicative ability of different viral strains by inoculating the strains in the culture at the same, typically low, MOI, and follow accumulation of viral genomes over time [47, 48]. However, if two viral strains differ in the degree of apparent cooperativity, the same dilution of the stock assuming linear reduction in MOI with dilution may in fact results in dramatically lower MOI for viruses with higher *n* – for example, 10 fold dilution of viral stock would result in nearly 40 fold reduction in infections for a strain with cooperativity degree of *n* = 1.6 (**Figure 6C**). Such a strain may then show delay in accumulation of viral particles in culture that may be incorrectly interpreted as reduced replication capacity and not reduced primary infection efficacy [47, 48].

## Discussion

A single hit model, in which one infectious viral particle can give rise to a productive infection of a cell, has been commonly assumed to be correct, excluding, of course, multipartite viruses whose segments are packaged in separate particles. In particular, Pradhan *et al.* [16] write that in accord with a single hit model, “Most mammalian viruses show a linear relationship between the number of plaques and dilution of virus plated, holding the framework true in titration assays to determine viral titers and the infectious dose.” In this study and in our analysis of data from others’ studies, we show that a linear relationship between the frequency of infected cells and the virion (or genome) concentration in a sample is an exception rather than the rule for at least three different viruses (HCMV, HIV, VV) and different strains of HCMV (TB, TR, ME) showing that frequency of infected cells often increases faster than linear with higher virion concentration (**Figure 2 and Supplemental Figure S6**). In our experiments the degree of such apparent viral cooperativity (characterized by a parameter *n*, see **eqn. (1)**) was dependent on specific stock preparation, HCMV strain and, surprisingly, on the type of target cells used (**Figure 3**). We developed stochastic simulations to test whether different hypotheses about how viruses infect target cells may result in apparent cooperativity (**Figure 1B**); the basic/null model that allows for variability in virion infectivity and cell resistance to infection did not result in apparent cooperativity (**Figure 4A**). Two alternative models — accrued damage and virus compensation models – consistently predicted deviation of the infection curve from the single hit model with *n >* 1 (**Figure 4C-D**), and a model in which virions form clumps did not always result in apparent cooperativity (e.g., **Figures 4 and 5** but see **Supplemental Figure S12**). By using DLS we showed that the distribution of clumps in our stock of HCMV-TB strain is skewed; we predict that most of these clumps consist of single virions but few clumps have multiple virions (**Supplemental Figure S11**); this is similar to a previous observation made for VV [33]. Presence of apparent cooperativity (*n >* 1) makes traditional calculations of the stock titer, and as a result, the MOI in a given experiment, imprecise, especially if the degree of apparent cooperativity varies between viral strain (**Figure 6**).

Our results are in line with a notion that outcome of infection is dependent both on the virus and target cell details and is highly stochastic at low virion to cell ratios [5, 24, 49–51]. It is notable, though, that despite the importance of both virus and cell properties for cell infection, reference sources and classical textbooks when characterizing viruses, e.g., by their specific infectivity, often do not include cell details used for virus titration. Several previous studies have documented deviations of cell infection rate from the single hit/null model but in most of such cases the frequency of infected cells (or the number of viral plaques) increased slower than linearly with virion concentration [9, 10] (and see **Supplemental Figure S7**). One notable exception is rapid reduction in the frequency of plaques formed by the multipartite Guaico Culex virus on C6/36 cells with dilution of the viral stock [52]; fits of the power-law model to these (digitized) data suggested a high degree of apparent cooperativity of *n* = 4.5. Apparent cooperativity may help explain large differences in estimated specific infectivity of viruses between different labs [53] including even determining genome concentrations in a sample [54]. Recent studies also suggest that some metrics of viral stock infectivity, e.g., tissue-culture infectious dose 50 (TCID_50_), may not be always robust to the way measurements are performed [55].

Variable degree of apparent cooperativity between different viral strains may impact interpretation of experimental results [56]. For example, in comparing replication capacity of different strains it is common to infect cells at low MOI to allow for amplification of small differences in viral replication rates. However, if strains differ in degree of apparent cooperativity, the assumed MOI could differ between strains substantially (**Figure 6**). Therefore, the difference in viral titers observed late in culture may not be due to differences in viral replication rates alone [47, 48]. For example, a recent study documented a lower concentration of drug-resistant transmitted/founder (**T/F**) HIV strains over 8 day culture as compared to drug-susceptible HIV strains [47]. Most of this differences primarily arose during the first 2 days of culture and may have resulted in differences in actual MOIs and not in replication rates of the T/F viruses. Indeed, visual regression analysis suggests similar replication rates (i.e., slopes) for two sets of T/F viruses from day 2 onward (see **Figure 3A** in [47]).

Our study has several limitations. First, we did not block potential secondary infection by progeny virions produced by the cells infected by the primary inoculum. It is possible that cells that had been infected with more virions would start virion production earlier causing faster than linear increase in the fraction of infected cells with increasing genomes/cell values [56]. However, HCMV has a long replication cycle with viral progeny release beginning about 3-4 days after infection [26, 38–40, 57, 58] that is consistent with our time course data (**Supplemental Figure S2**). Therefore, by measuring the frequency of infected cells at 3 days post virion inoculation we should not detect secondary infections.

Second, while we have proposed alternative mechanisms that may result in apparent cooperativity, at present we could not discriminate between these alternatives, in part, because the models lacked specifics – e.g., if virions interacting with a cell reduce its resistance to infection, what does it mean exactly [12]? If virions in a collection augment their infectivity (which may be expected for segmented viruses), how does that viral compensation actually work? Designing experiments that would discriminate between these alternatives would require focusing on a specific mechanism. For example, it may be that that the initiation of gene expression is difficult but is more efficient when there are more virions bringing in more tegument transactivators like pp72/ppUL35 [59]. Alternatively, it may be that there is a bona fide resistance mechanism at play here (e.g. “interferon”) that is antagonized by a viral tegument protein (like TRS1/IRS1 that acts against PKR and 2’5’OAS) [60]. Accrued damage model is also consistent with the idea that at higher genome/cell values, the inoculum itself (including cell and/or virion debris) may impact overall susceptibility of all cells in the well, for example, making them more susceptible to infection. It may be expected, though, that exposing cells to debris would increase cell resistance to infection; this would result in *n <* 1 that we did not observe at small genomes/cell values. Addressing these hypotheses is an area of future research that will require funding.

We would note, however, that our observation that the degree of apparent cooperativity *n* depends on the target cell type, i.e., we found lower *n* on epithelial cells as compared to fibroblasts (**Figure 3A**) argues against clumping as the primary mechanism of apparent cooperativity. Indeed, in stochastic simulations we found that changing target cell resistance to infection did shift the infection curve but did not impact estimate of *n* (results not shown). We believe that the contrast between cell types will be an approach to understand mechanism(s) of apparent cooperativity. One difference is entry pathways, the ECs involve endocytosis and endosome acidification whereas the fibroblasts do not. There are clearly different receptors involved also, although they are not well characterized. One recent report showed that the gH/gL/UL128-131 complex (aka, “pentamer”) is not just dispensable for entry into fibroblasts, but inhibitory [61]. They suggest that the pentamer might bind to a receptor on fibroblasts that activates a pathways that acts against viral early-immediate expression, It could be that in this situation, more virions are really helpful to overcome that block, whatever it is.

Third, we do not yet have an analytical model that would accurately describe the whole relationship between infections/cell and genomes/cell for all data; a simple extension of the power-law model allowing for saturation in the infection frequency at low and high genomes/cell values did not always fit the data accurately (**Supplemental Figure S4**). We also do not have a clear explanation of why infection frequency declines at high genomes/cell values for some strain-cell combinations (e.g., **Figure 2A, C, D, I, J**). Because we measured cell infection in live cells, increase in cell death at higher genomes/cell values may result in the decrease in the number of viable cells. Fourth, even though we measured distribution of clumps in our viral stock of HCMV-TB-GFP, we did not determine how many virions there are in clumps of different sizes. Sorting such clumps and using qPCR may be a way to directly measure this critical parameter impacting importance of viral clumping in determining degree of apparent cooperativity [41]. Fifth and finally, while we assume that at higher genomes/cell values several virions likely initiate the infection we had no ability to measure this directly. Our future studies will use either the same strain or different strains expressing GPF or mCherry to directly measure the rate of cell coinfection with two or more virions.

Our work opens avenues for future research. The key point is that in the presence of apparent cooperativity (*n >* 1), MOI calculated from the stock does not scale linearly with virion concentration; therefore, for viral strains with apparent cooperativitiy (*n >* 1) infectious titer may need to be determined for multiple values of genomes/cell and at different target cell concentrations for each virus-target cell combinations in question (**Figure** 6C and **Box 1**). It will be important to determine if segmented viruses such as influenza A virus (**IAV**) exhibit even higher degrees of cooperativity than HCMV, as one would expect viral compensation to be important for production of infection of cells by IAV. Whether virion aggregation (or clumping) is the key mechanism explaining apparent cooperativity also will have to be tested for different virus-target cell combinations. Further complicating matters is the possibility that virions may cooperate without the requirement to coinfect the same cell. Faba bean necrotic stunt virus (**FBNSV**), a multipartite plant virus, was shown to share gene products between neighboring infected plant cells, cooperating to provide viral proteins that would otherwise be lacking, due to infection by only a portion of the genome segments [62]. It would be interesting to explore if such cooperation would be detected by our methodology.

- Match viral titration methods to the experiment as far as possible. This includes using the same dilution of the viral stock, the cell type, duration of inoculation, and readout of infection.
- When possible, determine the degree of apparent cooperativity (“*n*”-value, **eqn. (1)**) for each virus strain/cell type pair being studied.
- If *n* = 1 (no cooperativity), it is reasonable to calculate experimental MOI based on stock infectivity value determined from a convenient stock dilution.
- If *n >* 1 or unknown, then stock infectivity should be determined at a dilution resulting in an MOI as close as possible to the desired experimental MOI. Alternatively, the inoculum size can be empirically determined to yield the desired number of infected cells. In these ways different virus/cell type pairs can be compared more fairly.

**Box 1**: Recommendations on titrating viral stocks and on performing experiments when comparing different viral strains.

Our work has focused on the initial steps of viral life cycle – infection of a cell. How changes in the virion concentration cells are exposed to impacts the number of virions produced per infected cell remains to be determined. Classical and more recent studies highlighted large heterogeneity in virion production by individual virus-infected cells for different viruses including IAV and HSV-1 [63]. In our preliminary experiments, we found that the number of HCMV-TB virions produced by fibroblasts was approximately independent of the genomes/cell used for infection, but more research is needed. Finally, there is a need to generate specific and testable hypotheses of how individual virions may cooperate to improve their chances to infect a cell. Manipulation with viral genes regulating entry of virions into the cell and/or efficiency of within-cell virus replication and with target cell genes regulating response to block virus replication would be highly instrumental. Development of more detailed mathematical models of within-cell virus replication may also generate novel hypotheses leading to apparent viral cooperativity [56]. Combining mathematical models and experiments to parameterize the models and test model predictions should help further understand how virions cooperate at infecting cells.

## Abbreviations

HCMV: human cytomegalovirus,
ECs: epithelial cells,
MOI: multiplicity of infection,
IU: infectious unit,
PFU: plaque-forming unit,
VV: vaccinia virus,
PDF: probability density function,
LRT: Likelihood ratio test,
DIPs: defective interfering particles,
DLS: dynamic light scattering.

## Data and code sharing

The data on HCMV infection from our experiments are available as a supplement to this paper and on github: https://github.com/Joshua-Miller-161/HCMV Dose Infection.git. Data on HIV infection of target cells from individual experiments (shown in Supplemental Figures S5A) were provided by Tenthorey *et al.* [64]. Data on infection of target cells by VV (shown in Supplemental Figures S5B) were digitized from the original paper [42]. Data on TMV infection of leaves was taken from the tables in Nourinejhad Zarghani *et al.* [10]. Codes (python and Mathematica) used to simulate cell infection are available on github (see link above).

## Ethics statement

No animal work.

## Author’s contributions

Original idea of the study came after talk BJR gave at the Microbiology department of the University of Tennessee. CP performed all experiments under supervision of BJR. VVG performed analyses of experimental data. JM performed most computational simulations under supervision of VVG with some contributions of VVG. VVG wrote the first draft of the paper and all authors read, edited, and agreed on the final version.

## Acknowledgments

This work was supported by the star-up funds from Texas Biomed and in part by NIH/NIAID grants R01AI158963 to VVG and R01AI097274 to BJR. Additional support was provided by the University of Montana (UM) Center for Biomolecular Structure and Dynamics (P30GM140963), the UM Center for Environmental Health Sciences (S10OD025019-01), and the UM Genomics Core and Montana INBRE Data Science Core, which are funded by the National Institute of General Medical Sciences (P20GM103474), the Office of the Vice President for Research and Creative Scholarship at the University of Montana, and the M. J. Murdock Charitable Trust.

## Supplemental Information

**Supplemental Table S1:** Standardized relationship between volume pipette dispenses and the actual weight of the dispensed volume (standard curve). We performed experiments by dispensing different volumes of the DMEM media (at 20^0^C room temperature) with a 1 ml serological pipette and measured their weights in three independent replicates (shown in different columns). The pipette had a standard error of 2% (denoted as ± values in the first column). Analysis of the data is shown in **Supplemental Figure S1A**.

| Volume (uL) | Sample 1 (g) | Sample 2 (g) | Sample 3 (g) |
| --- | --- | --- | --- |
| 1000 ± 20 | 0.9450 | 0.9468 | 0.9435 |
| 900 ± 18 | 0.8563 | 0.8401 | 0.8534 |
| 800 ± 16 | 0.7665 | 0.7677 | 0.7665 |
| 750 ± 15 | 0.7000 | 0.7115 | 0.7144 |
| 700 ± 14 | 0.6636 | 0.6765 | 0.6670 |
| 600 ± 12 | 0.5685 | 0.5700 | 0.5766 |
| 500 ± 10 | 0.4747 | 0.4747 | 0.4775 |
| 400 ± 8 | 0.3790 | 0.3842 | 0.3776 |
| 300 ± 6 | 0.2771 | 0.2825 | 0.2836 |
| 250 ± 5 | 0.2337 | 0.2327 | 0.2248 |
| 200 ± 4 | 0.1820 | 0.1850 | 0.1849 |
| 100 ± 2 | 0.0845 | 0.0865 | 0.0802 |

**Supplemental Figure S1:**
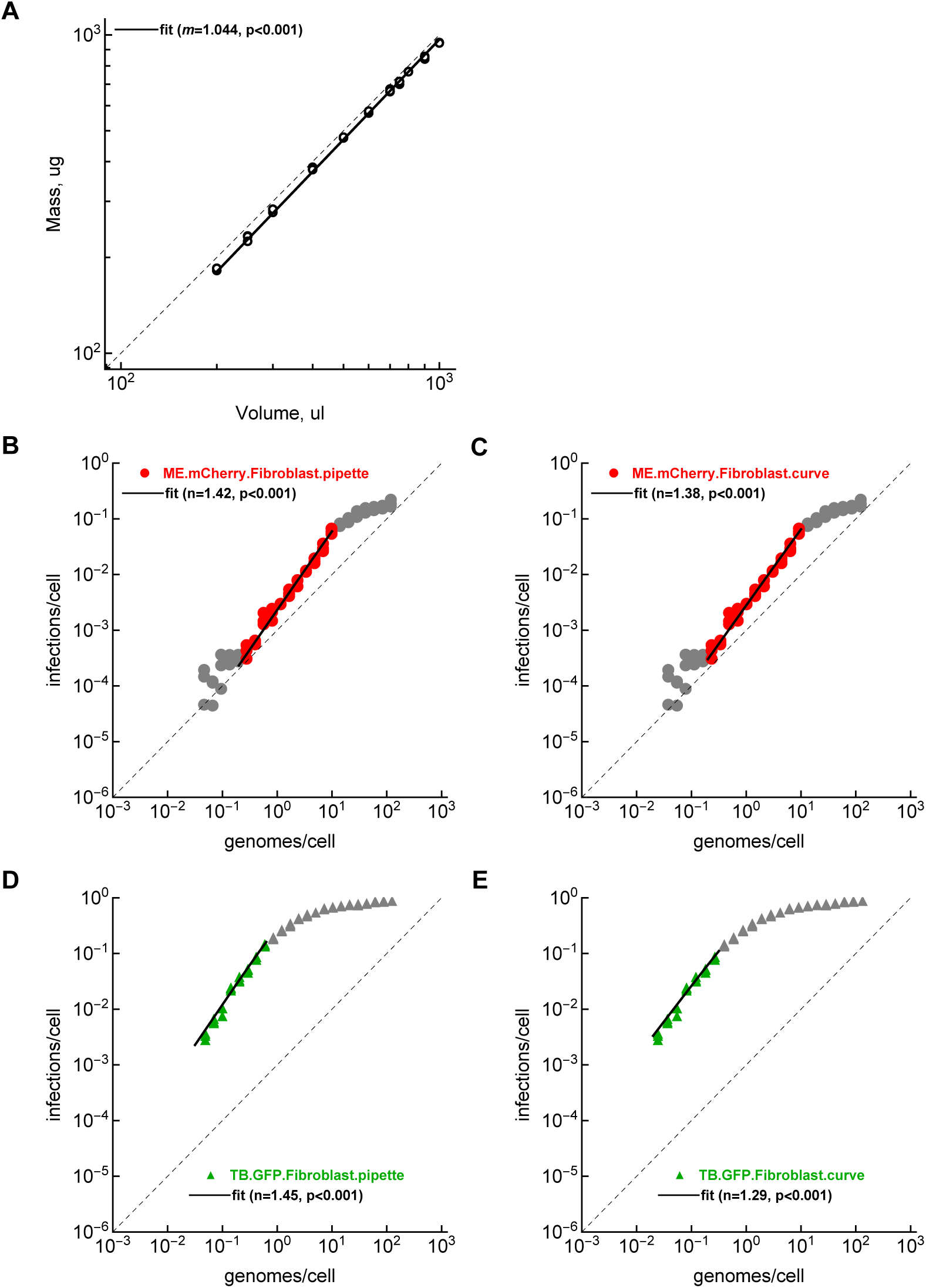
Using a standard curve converting media mass to volume allows for more accurate quantification of the degree of apparent cooperativity. **A**: We performed experiments in which we carefully measured the relationship between volume a pipette dispenses and the weight of this volume for a range of volume values in 3 independent samples (data are from **Supplemental Table S1**). Value *m* denotes the slope of the linear regression of log-transformed values and p-value is from the t-test for *m* = 1. **B-E**: We fitted the power-law model (**eqn. (1)**) to the data on ME-mCherry (**B-C**) or TB-GFP (**D-E**) strain infection of fibroblasts at different dilutions of viral stocks assuming that the pipette gives the correct assumed volume (**B**&**D**) or when correcting the dispensed volume using the calibration curve (**C**&**E**, see also Materials and methods for more detail). Estimated degree of apparent cooperativity *n* is shown on individual panels. Dilution factor used in calculating genomes/cell is 1.43 and 1.44 in panels **B** and **C**, respectively, and 1.43 and 1.48 in panels **D** and **E**, respectively.

**Supplemental Figure S2:**
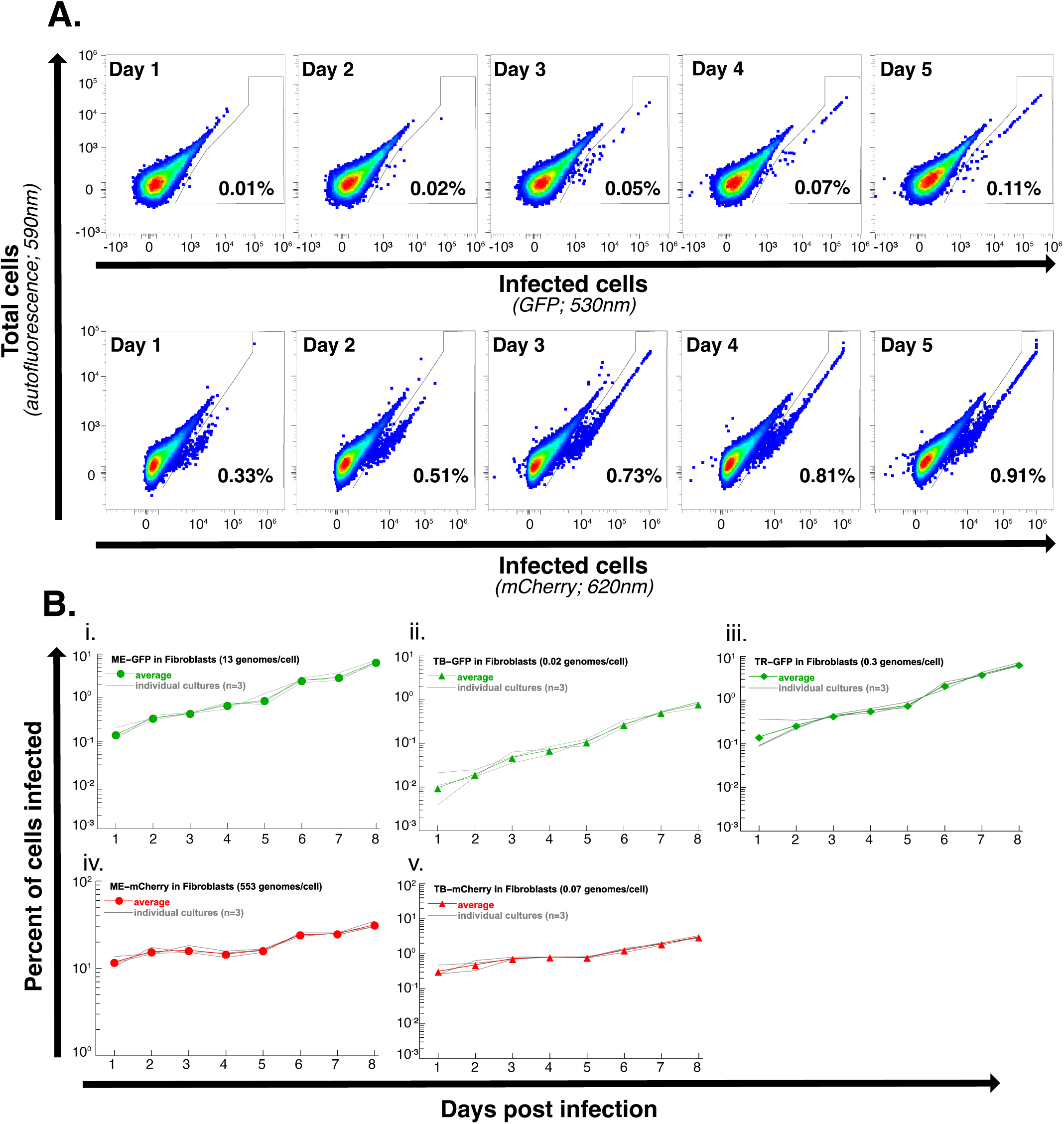
Kinetics of GFP and mCherry expression in fibroblasts exposed to different HCMV strains. **A**: We followed expression of GFP (top row) or mCherry (bottom row) in fibroblast over time after exposure to 0.02 genomes/cell of TB-GFP or 0.07 genomes/cell of TB-mCherry strains of HCMV. The frequency of cells detected as GFP+ or mCherry+ is shown on individual panels. **B**: We followed change in percent of fibroblasts infected with GFP- (**i-iii**) or mCherry-expressing viruses (**iv-v**). We show the data from individual three cultures (gray lines) and average (markers with colored lines).

**Supplemental Figure S3:**
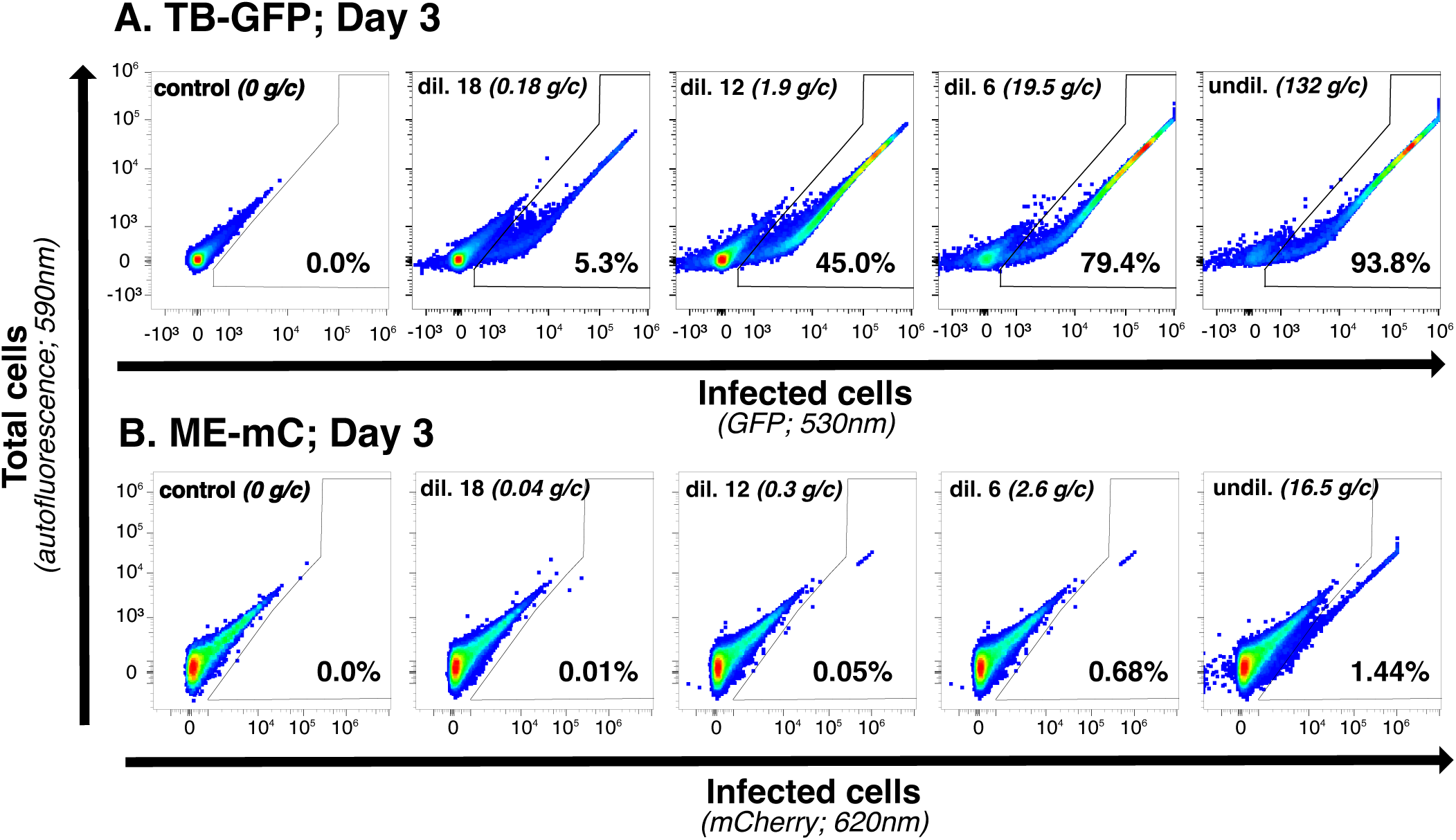
Detecting cells infected with GFP or mCherry-expressing HCMV virions at different dilutions of the stock. We show flow cytometry plots of TB-GFP- (**A**) or ME-mCherry- (**B**) exposed fibroblasts at different dilutions of the viral stocks detected at day 3 post exposure. Gates show the percent of cells detected as infected; specific dilutions and calculated genomes/cell (**g/c**) are shown on individual panels.

**Supplemental Table S2:** Estimates of parameters determining infectivity of HCMV strains for different target cells. The model fits are shown in **Figure 2** and details of how the model (**eqn. (1)**) was fit to data are given in Materials and methods. We list the HCMV strain (ME, TB, TR), marker expressed by the virus-infected cells (GFP or mCherry), and the type of target cells (Fibroblasts or Epithelial cells) used in experiments. Parameters are the infectivity of a single genome *p*(1) and the degree of apparent cooperativity *n*; numbers in parentheses are 95% confidence intervals generated by bootstrapping the data with replacement in 1000 simulations. Note that in **eqn. (1)**, *p*(1) = 1 − e*^−λ^*.

| Strain | Reporter | Cell type | $-\log_{10} p(1)$ | $n$ |
| --- | --- | --- | --- | --- |
| ME | GFP | Fibroblasts | 2.22 (2.24–2.21) | 1.34 (1.3–1.4) |
| ME | GFP | Epithelial cells | 4.35 (4.45–4.24) | 1.0 (0.88–1.09) |
| ME | mCherry | Fibroblasts | 2.52 (2.6–2.46) | 1.24 (1.18–1.36) |
| ME | mCherry | Epithelial cells | 3.92 (4.06–3.81) | 1.01 (0.91–1.15) |
| TB | GFP | Fibroblasts | 0.99 (1.02–0.96) | 1.61 (1.5–1.71) |
| TB | GFP | Epithelial cells | 2.66 (2.72–2.62) | 1.12 (1.09–1.16) |
| TB | mCherry | Fibroblasts | 1.59 (1.62–1.57) | 1.54 (1.47–1.61) |
| TB | mCherry | Epithelial cells | 3.73 (3.81–3.65) | 1.12 (1.06–1.17) |
| TR | GFP | Fibroblasts | 1.28 (1.31–1.25) | 1.27 (1.2–1.38) |
| TR | GFP | Epithelial cells | 3.17 (3.22–3.12) | 0.98 (0.9–1.18) |
| TR | mCherry | Fibroblasts | 3.39 (3.43–3.34) | 1.13 (1.04–1.2) |

**Supplemental Figure S4:**
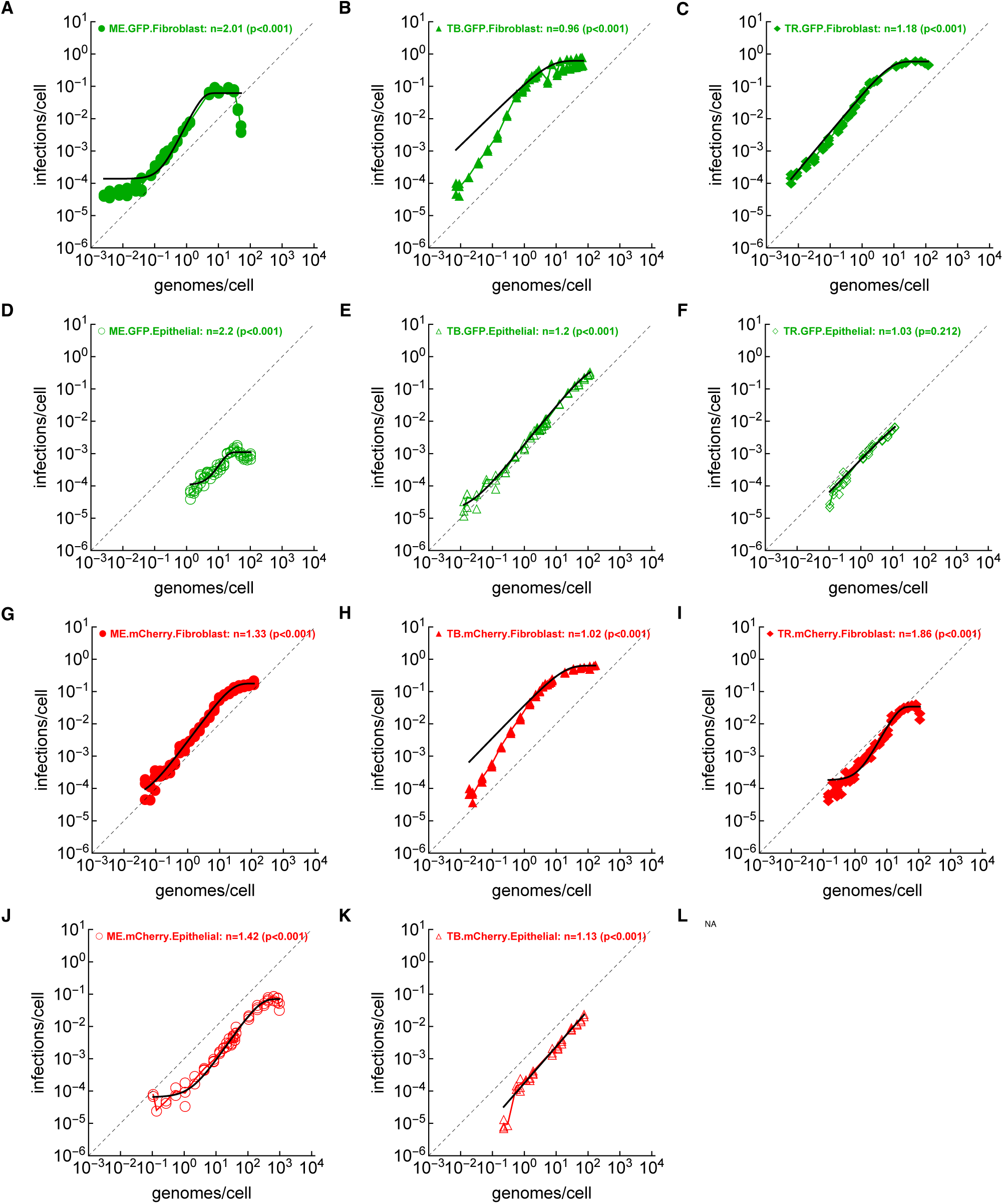
Apparent cooperativity of HCMV strains at infecting targets is still detected when fitting the extended power-law model to all data. We fitted extended power-law model (**eqn. (2)**) to the data; this model allows for saturation in infection probability at low or high genome/cell concentrations. Note that due to a large number of measurements at high genome/cell concentrations, the model fits are biased towards these datapoints in panels B and H.

**Supplemental Figure S5:**
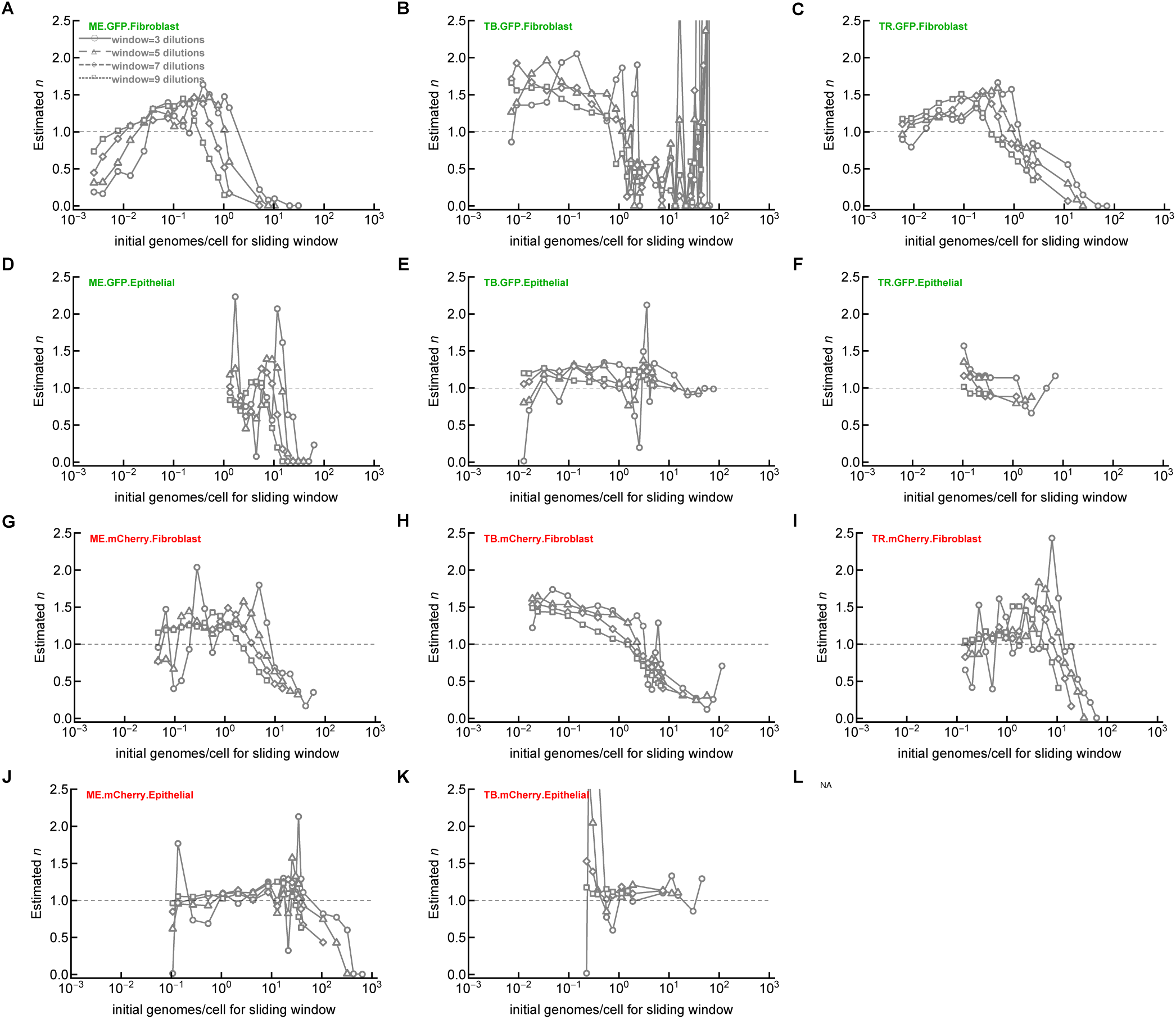
A degree of apparent cooperativity is dependent on the range of genome/cell and virus-cell combination. For each HCMV strain and cell combinations (see **Figure 2**) we determined the degree of apparent cooperativity *n* by taking subset of the data between different dilutions (starting with the highest dilution 23) and including different numbers of dilutions (denoted as *window*). We then fitted the power-law model (**eqn. (1)**) to these data and estimated the degree of apparent cooperativity *n*. Note that at high initial genome/cell subsets of data, *n <* 1 indicating competition between virions at infecting cells.

**Supplemental Figure S6:**
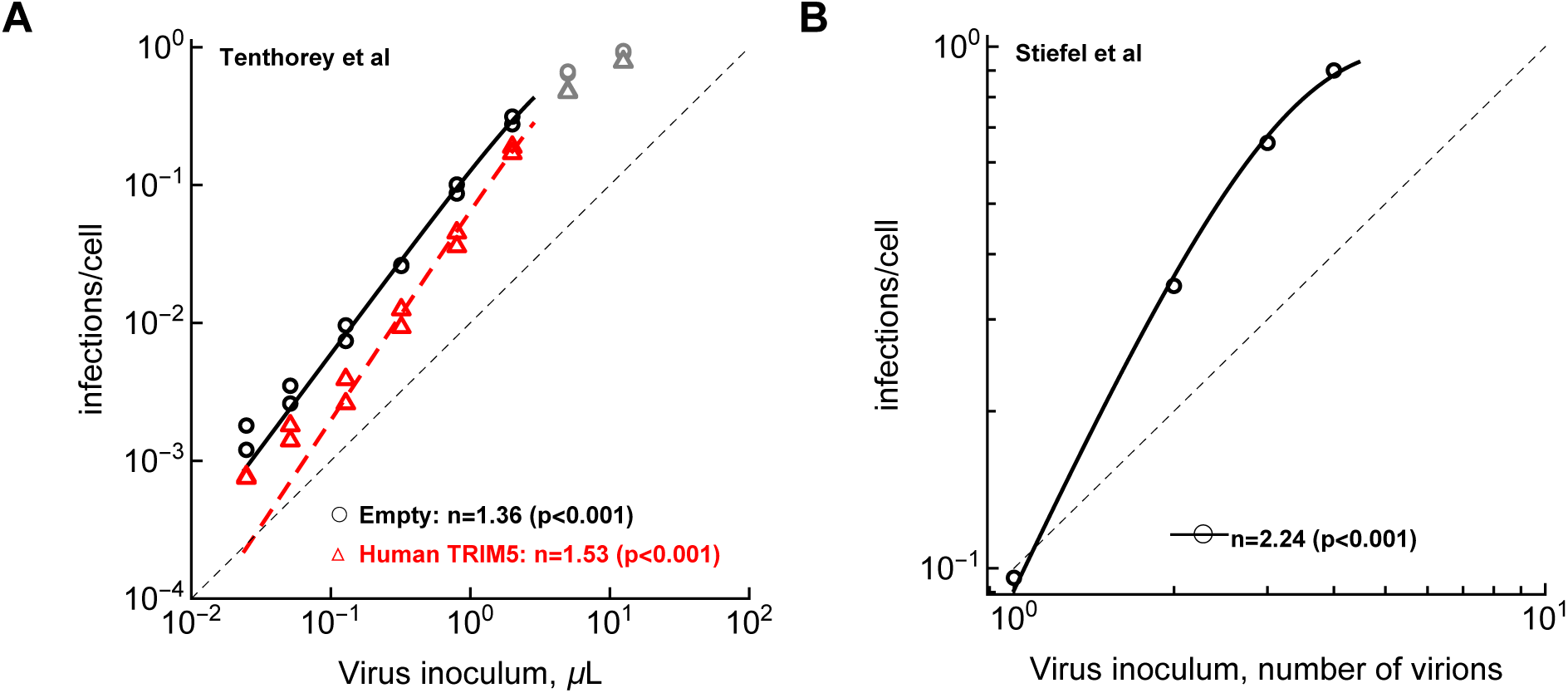
Apparent cooperativity of HIV and Vaccinia virus at infecting target cells. We analyzed data from two previously published papers [42, 64] that measured infection probability of cell exposed to different number of virions and estimated degree of apparent cooperativity from their data. **A**: change infection of TRIM5*α*-deficient CRFK cells (control or transdused with TRIM5*α*) exposed to different dilutions of HIV-1-GFP [64]. **B**: change in the probability of a HeLa cell exposed to a given number of vaccinia virus virions deposited on the cell surface [42]. For both studies we calculated the slope *n* by fitting the power-law model (**eqn. (1)**) to the data using maximum likelihood method (see Materials and methods for details). Shown *p* values are from likelihood ratio test when comparing model fit with *n* /= 1 to that with *n* = 1. Data are shown as markers and model fits as lines. Dashed line represents the slope of one. In **A**, data points shown in gray were excluded from the model fit.

**Supplemental Figure S7:**
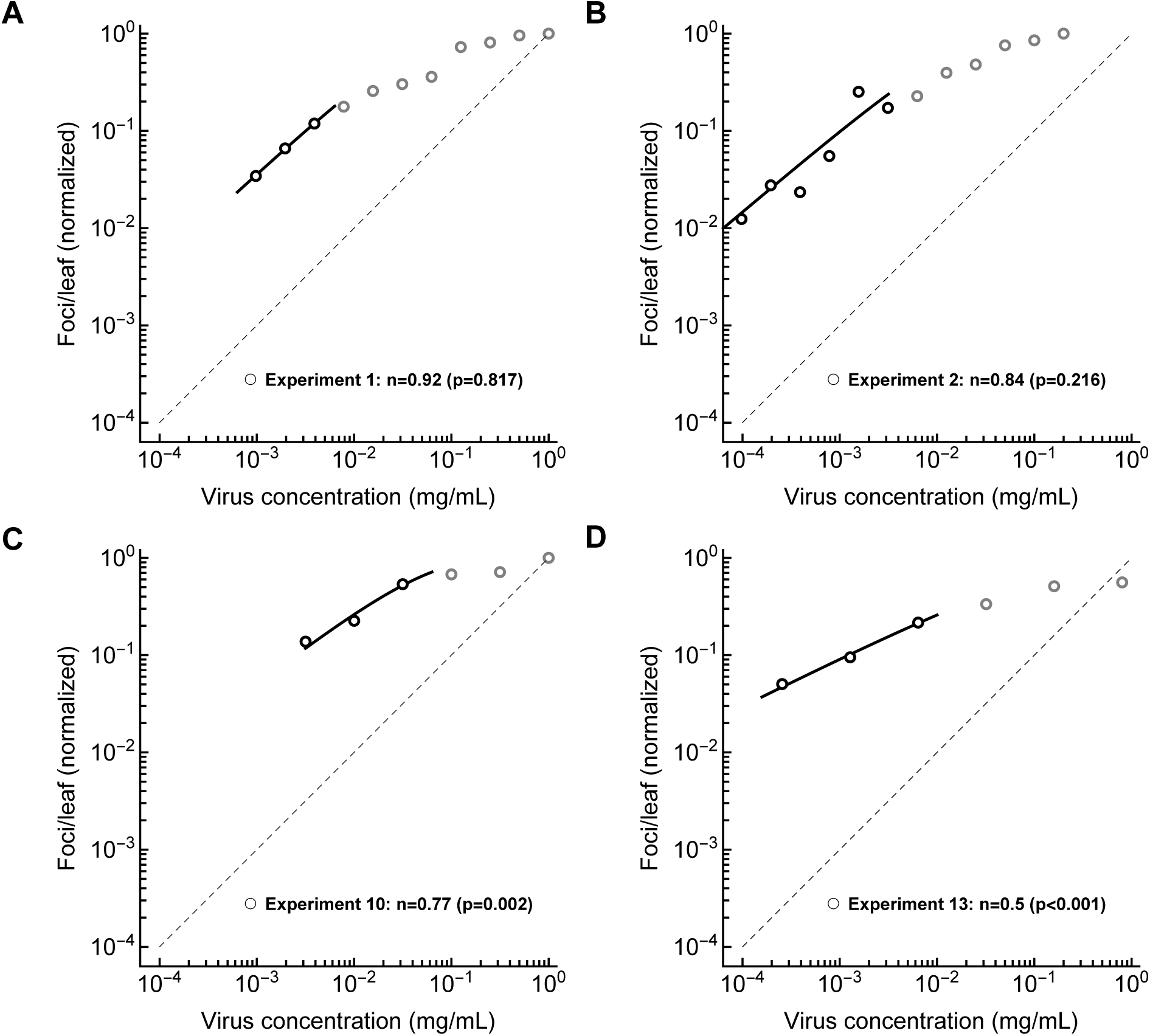
Lack of apparent cooperativity of Tobacco mosaic virus (TMV) on *Nicotiuna glutinom* plants. We analyzed data from studies documenting formation of lesions on plant leaves exposed to different dilutions of TMV stock [9, 10]. We show data that were numerically provided in Kleczkowski [9] and fitted the power-law model (**eqn. (1)**) to these data to estimate degree of apparent cooperativity *n* (listed p values are from a LRT with a model fit with *n* = 1). Dashed line represents the slope of one. Data points shown in gray were excluded from the model fit.

**Supplemental Figure S8:**
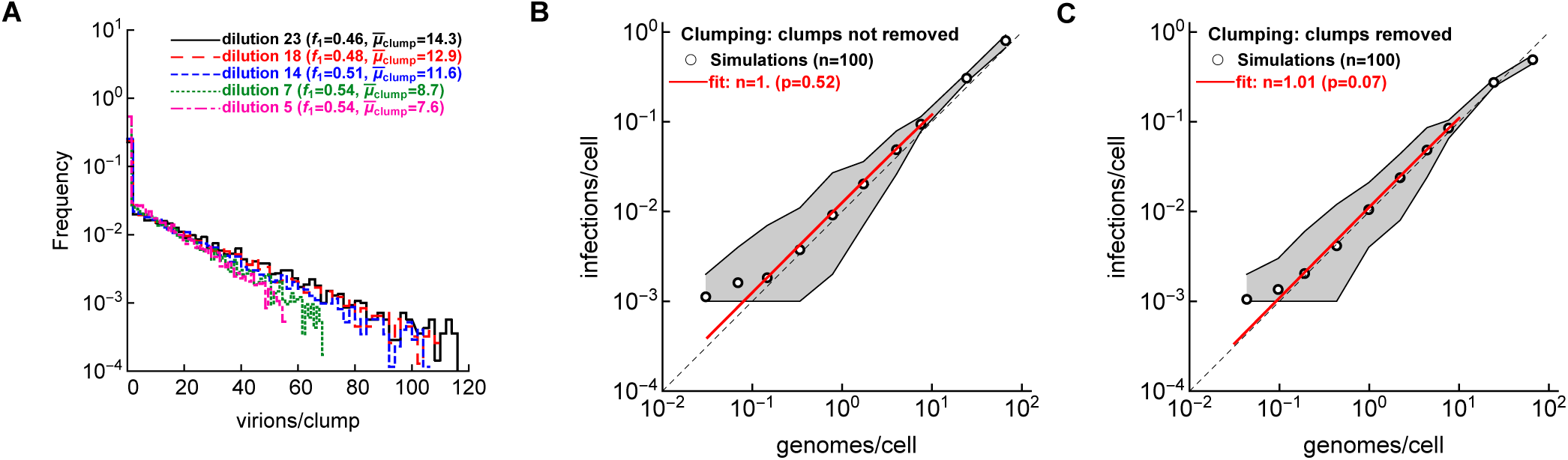
Removal of a clump from a well upon virion infection of a cell does not impact the estimated degree of apparent cooperativity in the clumping model. We simulated infection of cells by sampling virions in clumps (clumping model, **Figure 1**Bi) and calculating probability of cell infection at different genome/cell dilutions assuming that the distribution of virions per clump depends on the dilution (**A**). We run 100 simulations for each dilutions with parameters listed in **Table** 1 except 10^3^ cells per well, *R* = 15, *I* = 5, and we used geometric distribution to model the number of virions per clump. The fraction of clumps with a single virion (*f*_1_) and the average number of virions per clump (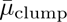) are shown for a selected numbers of stock dilutions in **A**. We show results of the simulations when clumps containing virions that successfully infected a cell are not removed from the well (**B**) or when such clumps are removed from the well upon successful infection (**C**). We performed 100 simulations per dilution (*n* = 100), and fit the power-law model (**eqn. (1)**) similarly to that shown in **Figure 4**. Other notations are the same as in **Figure 4**.

**Supplemental Figure S9:**
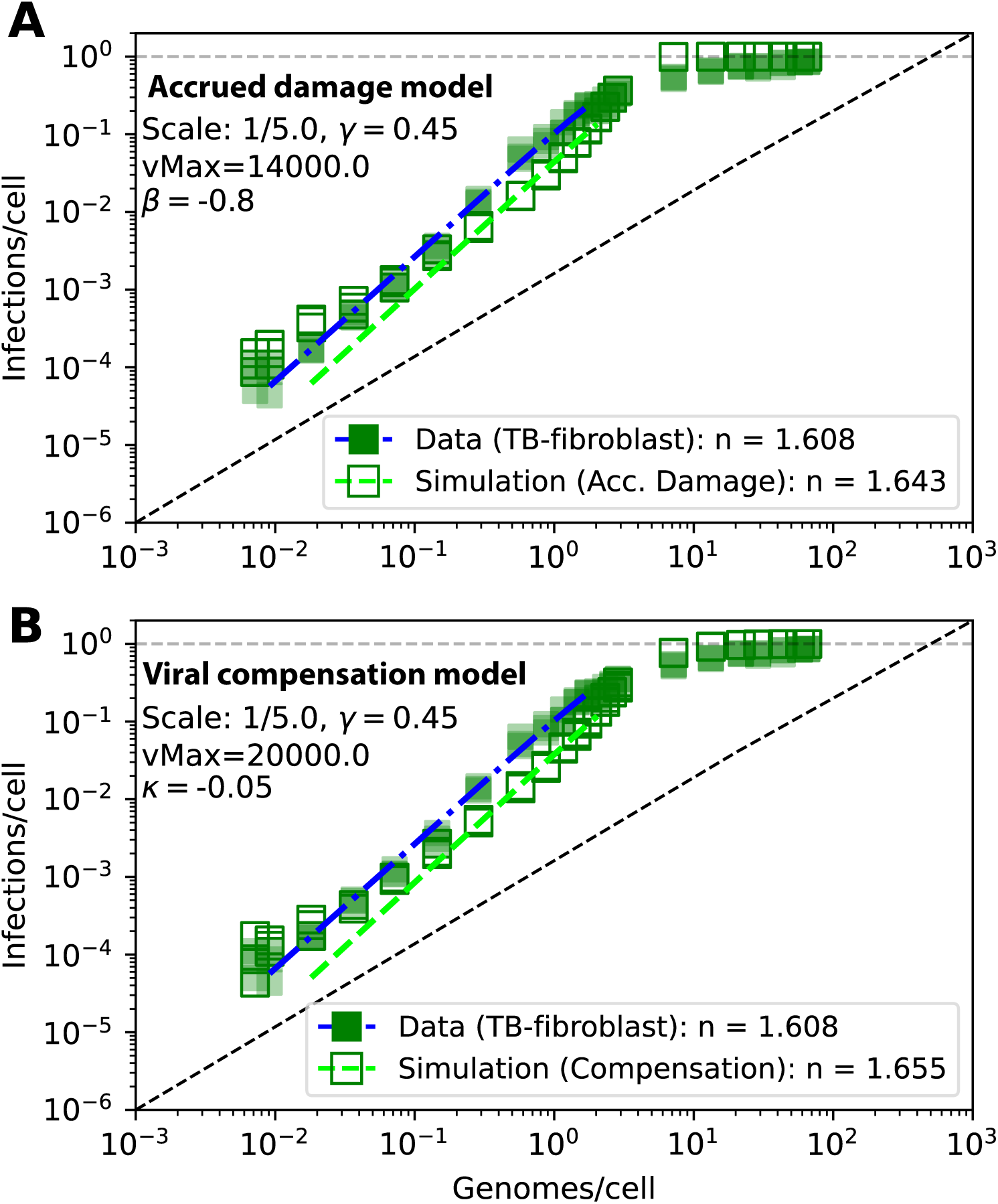
Both accrued damage and viral compensation models can reasonably match experimental data. We simulated infection of cells as in **Figure 4** but with slightly modified parameters (listed on individual panels) to more accurately match experimental data on infection of fibroblasts by HCMV-TB.

**Supplemental Figure S10:**
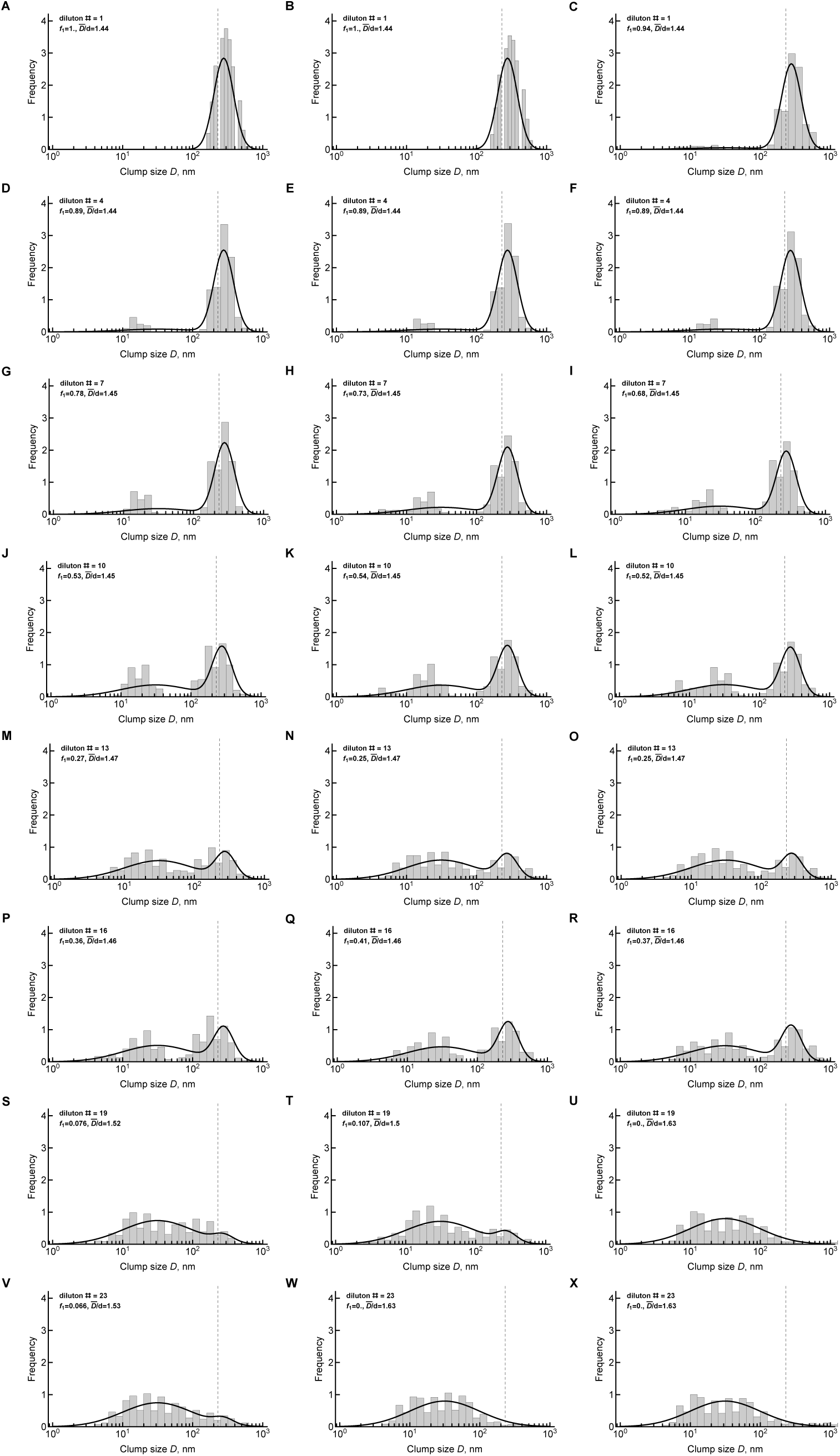
Distribution of viral clumps in three different experiments is well described by a mixture of two log-normal distributions. By using dynamic light scattering we measured distribution of viral clumps in three different preparations and for different dilutions of the viral stock of TB-GFP. We show measurements from three different viral stocks (in different columns). For each dilution we fitted a mixture of normal distributions to log-transformed clump size *D* and calculated the fraction of the larger peak *f*_1_; the average of the larger peak *D̅* is indicated on individual panels. A diameter of a single virion *d* = 230 nm is indicated by a vertical dashed line [34].

**Supplemental Figure S11:**
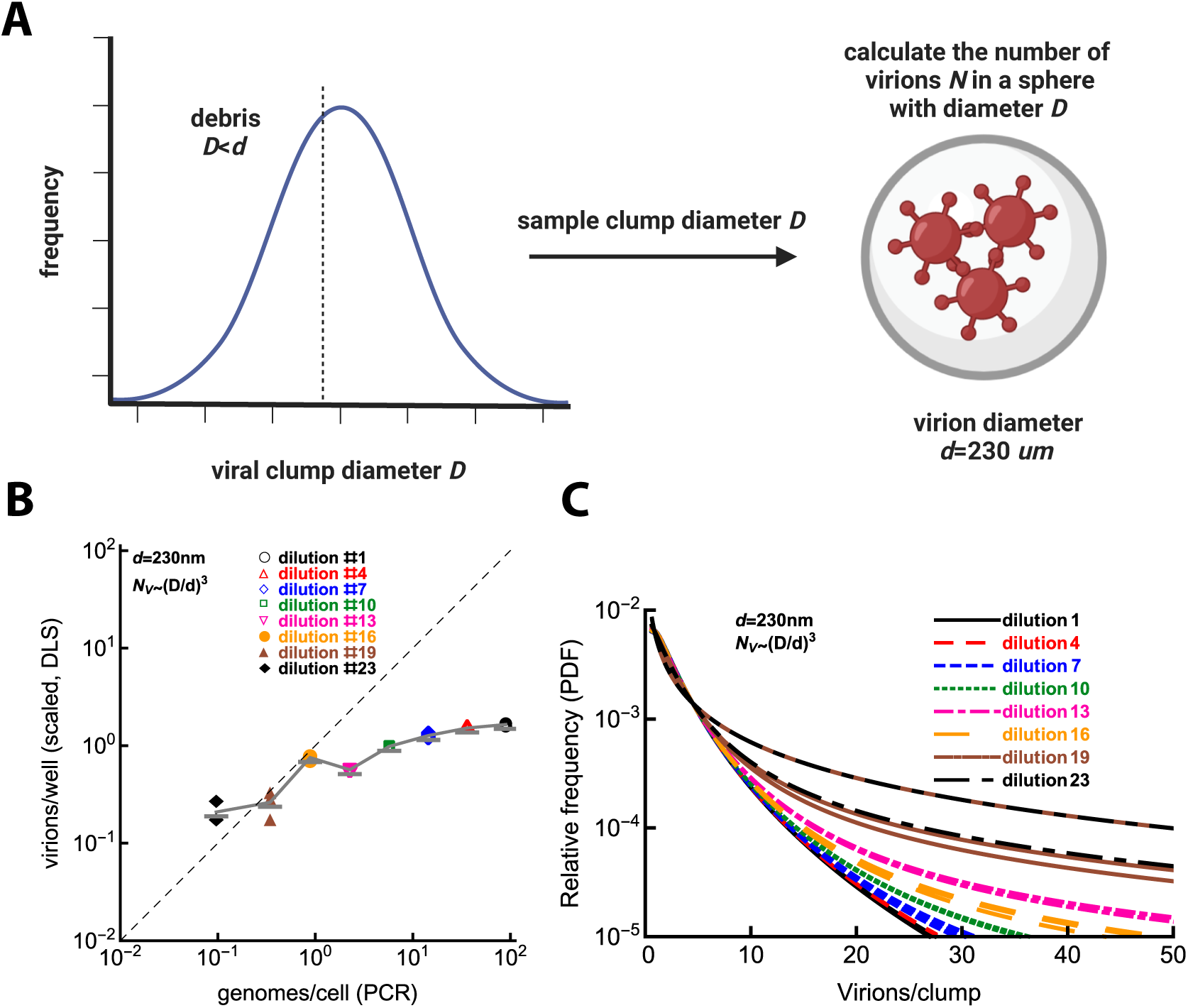
Quantification of virions in clumps does not result in linear scaling in virions/well at different dilutions of the stock. **A**: From the measured distribution of size (diameter) of viral clumps in a given stock preparation we sampled clumps of different sizes (with a diameter *D*). Assuming that these clumps are spheres and that a given clump is composed of spherical virions of a diameter *d*, we calculate the number of virions (genomes) per clump using **eqn. (5)**. Vertical dashed line in the plot denotes threshold for debris vs. virions when diameter of the clump is smaller than the virion size *d*. **B**: We calculated the predicted number of virions present in clumps of different sizes *D* using estimates of the log-normal distribution fitted to the data generated by dynamic light scattering (**Supplemental Figure S10**) and that number of virions per clump of size *D* scales as cube (**eqn. (5)**); the relative number of virions per well (or per cell) is proportional to 0.64 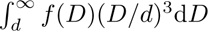. Each marker denotes individual stock dilution (columns in **Supplemental Figure S10**). Genome per cell were also calculated by PCR for this particular stock and a dilution factor 1.37. **C**: We plot normalized distribution of virions per clump assuming spherical packing and only accounting for clumps with *D > d* using model fits in **Supplemental Figure S10** for three samples (A, B, C) and 8 dilutions (24 curves in total).

**Supplemental Figure S12:**
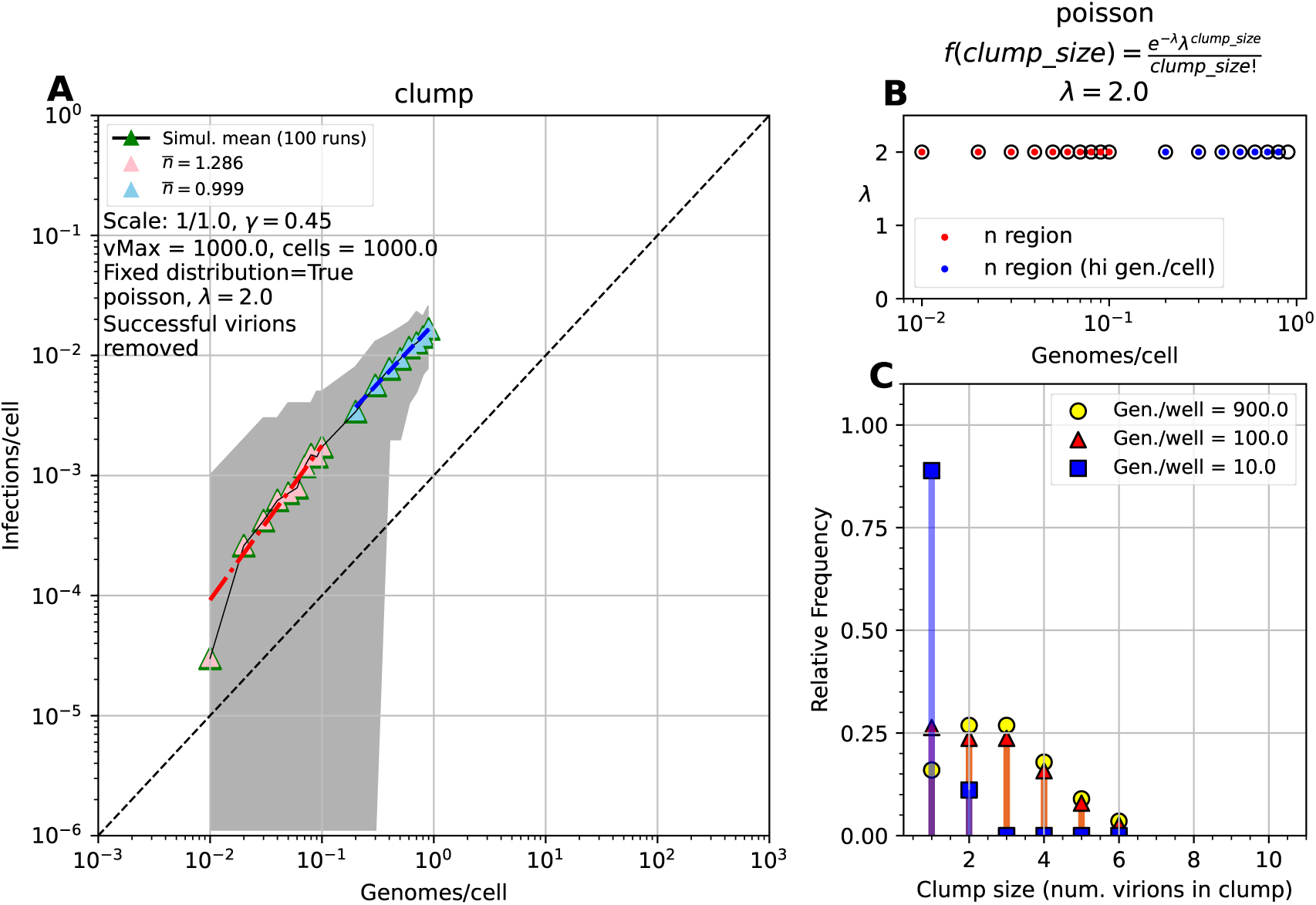
Clumping model may lead to apparent cooperativity at some chosen parameter values. We simulated infection of target cells by virions assuming that virions form clumps with the number of virions per clump following Poisson distribution (see Materials and methods for more detail). In these simulations we select subset of virions that would be titrated for a given well (and given dilution) and when subselecting virions attempting to infect a given cell we include all virions that belong to the clumps of the attempting-to-infect virions. Virions that succeed at infecting a cell are removed from a given well. **A**: We plot change in cell infection probability with increasing genomes/cell produced in simulations. We fitted the power-law model (**eqn. (1)**) to the subsets of simulation data and estimated degree of apparent cooperativity *n*. **B**: We show the mean of the clump size distribution *λ* at different genomes/cell values. **C**: We show an example of distribution of virions/clump simulated different stock dilutions. In these simulations we assume *λ* = 2 and we followed infection of 10^3^ cells/well; other parameters are also noted in **A**.

